# Normative trajectories of R_1_, R_2_* and magnetic susceptibility in basal ganglia on healthy ageing

**DOI:** 10.1101/2024.07.17.603922

**Authors:** Kwok-Shing Chan, Marcel P. Zwiers, Michelle G. Jansen, Joukje M. Oosterman, David G. Norris, Christian F. Beckmann, José P. Marques

## Abstract

Quantitative MRI techniques, including R_1_, R_2_*, and magnetic susceptibility mapping, have emerged as promising tools for generating surrogate imaging markers of brain tissue microstructure, enabling non-invasive in vivo measurements associated with myelination and iron deposition. Gaining insights into how these quantitative measurements evolve throughout a normal lifespan can enhance our understanding of brain maturation processes and facilitate the study of disease-related microstructural changes by distinguishing pathological alterations from normal brain development. In this study, we established the normative trajectories of R_1_, R_2_*, and magnetic susceptibility in the basal ganglia at 3T. We used a healthy ageing cohort comprising 260 subjects with an evenly distributed age range and sex ratio throughout adulthood. Utilizing the non-parametric Gaussian Process Regression model to derive the normative trajectories, we found that R_1_ in these structures predominantly exhibit a quadratic shape over age, while R_2_* and magnetic susceptibility are primarily linear. We validated the normative trajectories of R_2_* and magnetic susceptibility using an independent cohort. Additionally, we demonstrated that the spatial distributions of the quantitative MRI parameters also change with age in the putamen and caudate nucleus. This study not only reinforces existing findings on the association between age and qMRI but also provides valuable resources for studying cognitive ageing, in conjunction with the behavioural data available in the same data collection.

## 1. Introduction

Studying brain development across the lifespan is crucial for understanding brain maturation and establishing a standard framework to help us evaluate neurological disorder-induced deviations from typical quantitative phenotypes at an individual level. Traditional studies often involve group comparisons, such as between healthy individuals and patients, to distinguish pathological features from normal brain development, necessitating matching group demographic characteristics. Factors like age and sex are known to influence various brain (micro)structures (Gur et al., 2002; Jäncke et al., 2015; Persson et al., 2015; Xu et al., 2000). However, simply matching demographic characteristics may not be sufficient, as this overlooks how these factors might influence the effect of interest. The development of normative models allows for the exploration of deviations related to diseases from what is considered normal (Marquand et al., 2016a; Rutherford et al., 2023), considering their dependence on these factors. This approach transcends classical case-control designs, allowing researchers to better comprehend disease progression and the effects of medical interventions, without the need for acquiring prohibitively large control datasets due to cost and logistical constraints.

Most of the existing literature on normative MRI data focuses primarily on changes in brain structure, such as variations in tissue volume and cortical thickness (Bethlehem et al., 2022; Ramanoël et al., 2018; Scahill et al., 2003; Terribilli et al., 2011). This emphasis on morphometric characteristics stems from the relative nature of the underlying MR signal rendering the intensity of imaging voxels arbitrary, and the growing availability of conventional structural images across various large public datasets (Bookheimer et al., 2019; Jack et al., 2008; Miller et al., 2016; Satterthwaite et al., 2016; Snoek et al., 2021). Prior investigations have indicated a rapid increase in cortical thickness and tissue volumes during the early stages of development, followed by a gradual decrease after 1-2 years for cortical thickness and 18 years for total tissue volume (Bethlehem et al., 2022). Furthermore, studies have observed morphological disparities related to sex and hemispheric asymmetry (Bethlehem et al., 2022; Park et al., 2004; Taki et al., 2011; Watkins et al., 2001). In addition to cortical thickness, the framework of normative modelling can be expanded to encompass various other imaging-derived phenotypes, including diffusion-derived metrics in white matter among the most prevalent, and quantitative MRI (qMRI) properties.

Quantitative MRI parameters, including the longitudinal relaxation rate R_1_, effective transverse relaxation rate R_2_* and tissue magnetic susceptibility *χ*, are gaining prominence as surrogate biomarkers for myelin (Lutti et al., 2014; Rooney et al., 2007) and iron (Ghadery et al., 2015; Langkammer et al., 2012; Pontillo et al., 2022; Stüber et al., 2014) concentrations, respectively. Quantitative MRI offers direct and quantitative assessments of tissue MR properties. This approach not only holds the promise of furnishing more precise information about brain tissue composition but also of generating metrics that are reproducible across sessions and scanners (Deh et al., 2015; Körzdörfer et al., 2019; Leutritz et al., 2020; Lin et al., 2015; Teixeira et al., 2019). These methods are becoming more readily available in MRI scanners and cohort datasets. Their reproducibility and sensitivity have prompted various studies demonstrating that R_1_, R_2_* and *χ* vary throughout the lifespan (Kühne et al., 2021; Li et al., 2023, 2014; Miletić et al., 2022), with these changes being specific to regions of interest (ROI). In traditional age-matched case-control studies, these metrics are significantly affected in various brain regions depending on the diseases, reflecting, for example, increased iron deposition in the putamen and substantia nigra in Alzheimer’s and Parkinson’s diseases (Acosta-Cabronero et al., 2013; Pyatigorskaya et al., 2020), respectively.

Summary statistics, like mean or median, are widely used to study changes in qMRI through ROI-based analysis (Seiler et al., 2020). Previous research has outlined normative trajectories in adulthood, showing an inverted U-shape for R_1_ (Erramuzpe et al., 2020; Yeatman et al., 2014) and linearity for R_2_* and *χ* (Acosta-Cabronero et al., 2016; Burgetova et al., 2021; Keuken et al., 2017; Li et al., 2021). Notably, recent findings suggest that both ageing and diseases can impact the spatial distribution of these quantitative parameters (Drori et al., 2022; Oldehinkel et al., 2022) and manifested in higher-order statistical analysis (Li et al., 2019). Alternatively, normative modelling of qMRI can be conducted at the voxel level, as exemplified in the work of Piredda et al. (Piredda et al., 2023, 2020). These quantitative atlases serve as valuable resources for facilitating automated pathology detection at the voxel level. However, this approach mandates high-quality non-linear image registration to align individual data with the atlases, a task compounded by individual-level morphological differences, especially in cortical grey matter studies, where structural variability is increased. For instance, the primary auditory cortex can consist of 1 or 2 Heschl’s gyri (Costa et al., 2011). Moreover, factors such as measurement noise and imaging artefacts can contribute to the derivation of the normative data, potentially reducing their sensitivity in detecting subtle changes.

In this work, we aimed to explore both the cross-sectional median and spatial effects of age on qMRI parameters R_1_, R_2_* and *χ* in the basal ganglia at 3T, utilising a healthy ageing cohort characterised by a broad age distribution. Our focus on subcortical nuclei stems from their robustness in *χ* measurements and the well-established association with iron accumulation. Our objectives were threefold: (1) to delineate normative trajectories across subcortical grey matter nuclei throughout the lifespan using a Gaussian Process Regression model along with conventional volumetric analysis; (2) to investigate the independence of deviations from normative values within each subcortical nucleus, as measured by different qMRI parameters; and (3) to examine the independence of variations from normative values across the subcortical nuclei, aiming to elucidate networks of co-variation from the norm. Lastly, we tested the utility of the normative trajectories in a validation dataset.

## 2. Material and Methods

### 2.1. Data acquisition

Data utilised to derive the normative trajectories is part of the Advanced Brain Imaging on Ageing and Memory (ABRIM) data collection (Jansen et al., 2024) available at https://doi.org/10.34973/7q0a-vj19. A complete description of the MRI protocol and the pre-processing steps can be found in (Jansen et al., 2024). Briefly, data acquisition was performed at 3T (Siemens, Erlangen, Germany) on 301 healthy volunteers. Forty-one participants were excluded due to incidental findings and/or severe motion artefacts in the data, resulting in data of 260 participants to be included in the statistical analysis (18-79 years; mean ± SD=50.8±16.8 years, see Table 1). Informed consents were obtained from all participants for being included in this study. The imaging protocol of the data analysed here consisted of:

1. Whole-brain T_1_ (=1/R_1_) scan using MP2RAGE (Marques et al., 2010), α_1_/α_2_ = 6°/5°, TI_1_/TI_2_=700 ms/2000 ms, TR/TE = 6000 ms/2.34 ms, resolution = 1 mm iso., acquisition time (TA)=7 mins;
2. Whole-brain turbo FLASH B_1_ mapping, TR/TE=10000/2.23, α_SAT_/α=80°/8°, resolution 3.3 mm x 3.3 mm x 2.5 mm, TA=20 s;
3. Bipolar 3D multi-echo GRE, TR/TE1/ΔTE = 44 ms / 6.14 ms / 4 ms, 9 echoes, GRAPPA = 3, phase/slice partial Fourier = 0.875/0.875, α=20°, resolution=0.8 mm iso., TA=9.5 mins.

**Table 1:** Demographics of the ABRIM dataset used in this work.

| ABRIM | Full range | Age range (years) |  |  |  |  |  |
| --- | --- | --- | --- | --- | --- | --- | --- |
|  |  | 18-30 | 31-40 | 41-50 | 51-60 | 61-70 | 71-79 |
| N | 260 | 39 | 41 | 44 | 49 | 43 | 44 |
| Age mean±SD (years) | 50.8±16.8 | 24.6±3.9 | 35.4±3.0 | 46.0±2.9 | 55.4±2.9 | 65.2±3.0 | 74.0±2.5 |
| N<sub>Male</sub> | 119 | 17 | 17 | 21 | 24 | 22 | 18 |
| Age mean±SD (years) | 51.2±16.2 | 26.3±3.1 | 34.3±3.1 | 46.0±3.2 | 55.4±2.8 | 65.4±3.0 | 73.7±2.6 |
| N<sub>Female</sub> | 141 | 22 | 24 | 23 | 25 | 21 | 26 |
| Age mean±SD (years) | 50.5±17.4 | 23.4±4.1 | 36.1±2.8 | 46.0±2.7 | 55.5±3.0 | 65±3.0 | 74.1±2.6 |

To study the applicability of the R_2_* and *χ* normative models derived from the ABRIM datasets, an independent cohort with a similar imaging protocol, which is part of the “Parkinson Op Maat” (POM) data collection (Bloem et al., 2019), was used. Briefly, the POM study followed 500-550 patients who were diagnosed with Parkinson’s disease for 2 years aiming to gain insight into the onset and course of Parkinson’s disease. During the study, two MRI scans were made for each participant (within 5 years of first diagnosis and 2 years after the first scan). Here, we only included the first scan of the patient data and either the first or second scan of the healthy control data, depending on the data quality, for data analysis. In this analysis, data from 44 healthy volunteers and 316 patients were used. Data acquisition was performed at 3T (Siemens, Erlangen, Germany). The demographics are shown in Table 2. The imaging protocol of the data analysed here consisted of:

1. Whole-brain T_1_-weighted anatomical scan using MPRAGE, α=8°, TI=880 ms, TR/TE=2000 ms/2.03 ms, resolution=1 mm iso., TA=5 mins;
2. Bipolar 3D multi-echo GRE, TR/TE1/ΔTE=44 ms/6.14 ms/4 ms, 9 echoes, GRAPPA=3, phase/slice partial Fourier=0.875/0.75, α=20°, resolution=0.8 mm iso., TA=8 mins.

**Table 2:** Demographics of the POM datasets used in this work.

| POM | Control | Patient |
| --- | --- | --- |
| N (Male; Female) | 44 (20; 24) | 316 (175; 141) |
| Age mean±SD (years) | 58.2±9.2 (60.3±8.7; 56.4±9.5) | 60.9±9.0 (60.8±9.1; 60.9±8.9) |

### 2.2. Data Analysis

#### 2.2.1. Data processing

SynthStrip (Hoopes et al., 2022) was used for skull stripping on the second inversion time of the R_1_ data (INV2, proton density contrast) with the ABRIM cohort or the MPRAGE image with the POM cohort, and the first echo of the GRE data (GRE1). For *χ* processing, the GRE brain masks were first refined by excluding voxels with high R_2_* values on the brain mask edge to improve estimation robustness. *χ* maps were derived using SEPIA v1.2.2.4 (Chan and Marques, 2021) with the following pipeline: bipolar gradient phase correction (Li et al., 2015), ROMEO (Dymerska et al., 2021) for total field computation, V-SHARP (Li et al., 2011) for background field removal and LP-CNN (Lai et al., 2020) for dipole field inversion. A retrospective correction method was implemented to correct those datasets acquired with a tiled field-of-view with respect to the main magnetic field direction before the dipole field inversion step (Kiersnowski et al., 2023). The mean susceptibility value across the whole brain was used as a reference. For R_2_* mapping, Marchenko-Pastur Principal Component Analysis (MP-PCA) denoising (Veraart et al., 2016) with a patch size of 3×3×3 voxels was applied to the complex-valued mGRE data to improve SNR and the magnitude of the denoised data was then extracted to derive R_2_* maps based on a closed-form solution (Gil et al., 2016).

Quantitative R_1_ maps were obtained using the BIDS-compatible Matlab function *“bids_T1B1correct.m*” hosted on GitHub (https://github.com/Donders-Institute/MP2RAGE-related-scripts). This function performs (1) registration of the magnitude image from the turbo FLASH B_1_ map to the MP2RAGE image with INV2 using SPM12 (Penny et al., 2006); (2) the resulting transform matrix was used to register a spatially smoothed version of the B_1_^+^ map to the MP2RAGE space, which was then used to (3) correct for the B_1_ transmit inhomogeneities in R_1_ estimations (and derive M_0_ maps over a wide range of R_1_ values). In the last step, a fingerprinting-like approach (Ma et al., 2013) was used to estimate R_1_ instead of the traditional MP2RAGE lookup table (Marques et al., 2010). This was achieved by searching the maximum likelihood of the inner product of the signal at the two inversion times by a dictionary of simulated signals across a predefined range of R_1_ and M_0_ values. The MP2RAGE signal dictionaries were generated with a step size of 0.005 nominal B_1_ field.

#### 2.2.2. Subcortical grey matter parcellation

All data and statistical analyses were conducted in the common MNI space (ICBM 2009c Nonlinear Asymmetric) (Fonov et al., 2009). The entire image registration procedures are shown in Figure 1a. The MuSus-100 atlas (He et al., 2022) was used to provide parcellation labels of the basal ganglia, including detailed thalamic sub-regions. Co-registration between R_1_ and GRE data was achieved using a rigid body transformation between the skull-stripped MP2RAGE’s INV2 image and the GRE1 image. This transformation matrix was used to transform the *χ* and R_2_* maps into the R_1_ space for each subject. To facilitate high-quality subcortical grey matter parcellation, a 4-step registration procedure was performed as explained as follows.

**Figure 1:**
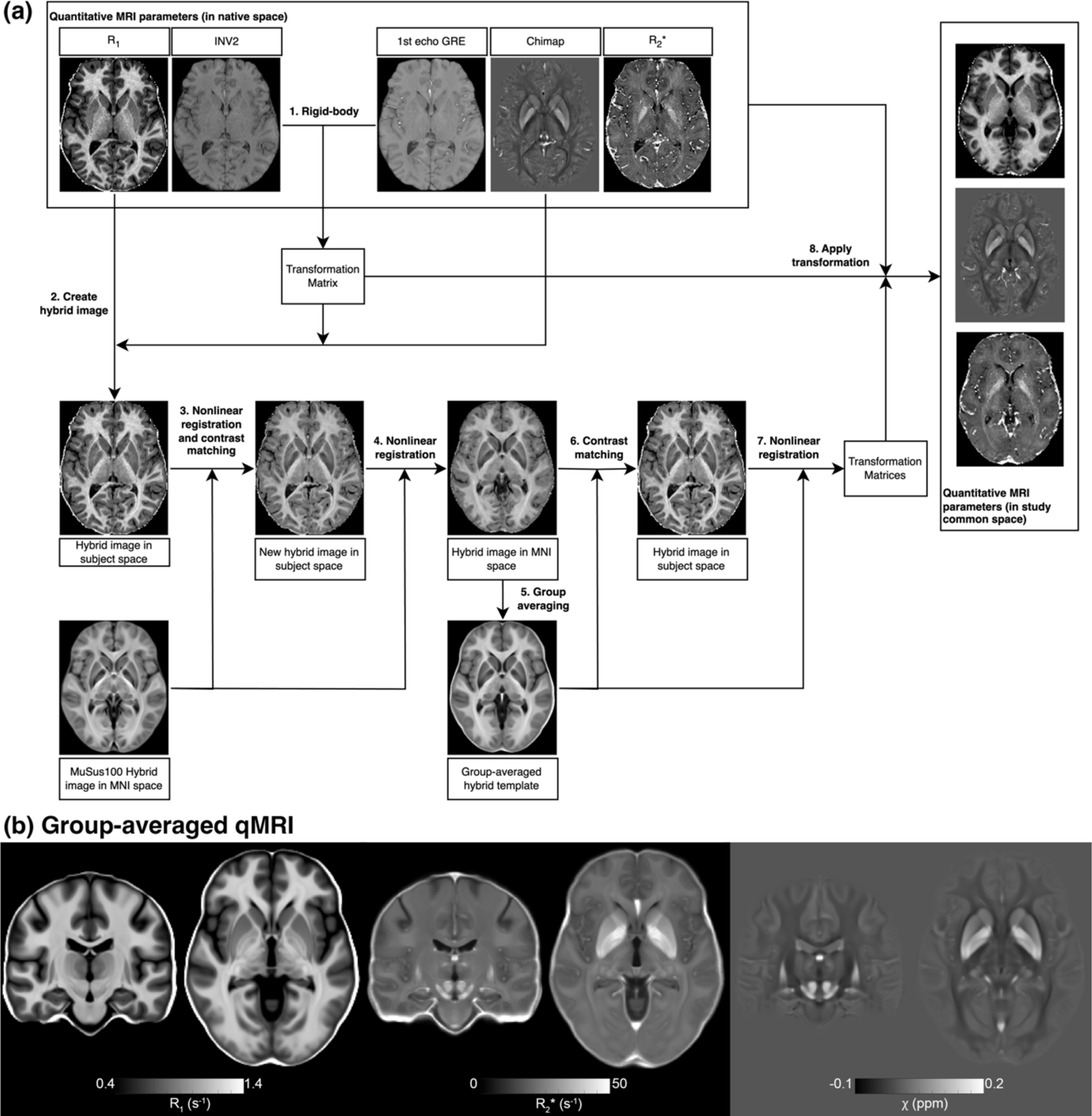
(a) An illustration of the image registration process on the three qMRI parameters maps. (b) Example R1, R2* and *X* maps derived by averaging data across all subjects in the study common space.

The first step involved the creation of an initial R_1_-*χ* hybrid image, which was then registered to the MuSus-100 atlas based on nonlinear transformation (Avants et al., 2011). The hybrid image was created using the following equation (Zhang et al., 2018):

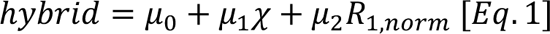

where R_1,norm_ is the R_1_ map normalised to values between 0 and 255 (clipping 1% brain masked values on each side of the histogram). μ_0_, μ_1_and μ_2_ were set to 0, 400 and 1 according to (Zhang et al., 2018). Since the hybrid image was derived from the R_1_ map here instead of the T_1_-weighted image of MPRAGE and a different algorithm was used to derive the *χ* map, the contrast of the initial hybrid image does not match exactly the hybrid template of the atlas but the improved contrasts between brain tissues do provide a fair alignment between the subject’s and the atlas data. This procedure has now been integrated into the SEPIA toolbox as of version 1.2.2.4.

The second step involved creating a hybrid image closely resembling the atlas template. To achieve this, we performed image contrast matching to enhance the similarity in image intensity between the subject’s R_1_ map and the atlas’s T_1_w template. This was performed by fitting a second-order polynomial function to the median value of all ROIs in the brain based on the parcellation labels of the atlas. A similar procedure was performed on the *χ* map, but in this case, a first-order polynomial function was used. Subsequently, the polynomial coefficients in Eq. (1) were derived from the atlas’s hybrid template with the matched subject’s R_1_ and *χ* data, which were then used to compute a new hybrid image for nonlinear registration. This step is crucial as it significantly improves the delineation of subcortical grey matter structures, as demonstrated in Supplementary Figure S1.

In the third step, a group hybrid template was created by averaging the hybrid images from all subjects. We then repeated the procedures in the second step with the new group-averaged hybrid image instead of the atlas’s template, as the new group-averaged image contains the same tissue contrast as our cohort data. All the qMRI maps were transformed into the group template space using the registration results of this step. The first three steps ensure high-quality image registration for all data to be transformed into a common space.

As a final step, subcortical grey matter structure parcellation was performed by using a nonlinear image transformation to register the atlas hybrid template to the group hybrid template (Figure 1a). For the POM cohort, the same registration procedures were repeated within the cohort datasets to obtain the subcortical grey matter parcellation, with the only change being using the T_1_-weighted MPRAGE image instead of the R_1_ map for image registration.

#### 2.2.3. Metrics of interest

To further minimise the variations caused by imperfect image registration and potential outliers from the veins in computing the normal models, the median qMRI (R_1_, R_2_* and *χ*) values were calculated for each subcortical grey matter structure. In addition to analysing each thalamic sub-nucleus individually, we also process the thalamus as a whole structure by combining all thalamus-associated labels from the MuSus-100 atlas. Besides the median statistics, the interquartile range (IQR) and skewness of the distributions were computed on each subject.

The tissue volumes of all subcortical ROIs were also extracted and normalised by the intracranial brain volume. To facilitate visualisation of the evolution of different volumes on the same plots, the volume of a given ROI was normalised by the population mean (*V*_ROI_) and the percentage change in volume (*V*_ROI,%_) is studied as:

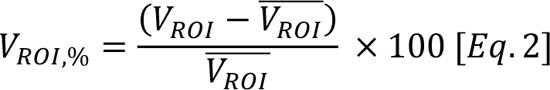

where *V*_ROI_ is the volume normalised by the intracranial brain volume.

To study the association between age and qMRI spatial distribution on the putamen and caudate nucleus, a first-order 3D polynomial function was employed to fit the spatial distribution across the ROI mask, which was eroded by 1 voxel in all directions to reduce the partial volume effect at the edges of the structures, so that the spatial gradients in the lateral-medial (L-M), posterior-anterior (P-A) and ventral-dorsal (V-D) directions of the MNI coordinate space can also be investigated in these structures, which can be expressed as follow:

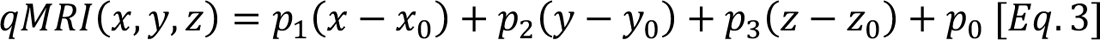

where *p*_1_, *p*_2_ and *p*_3_ describe the spatial gradient in the L-M, P-A, and V-D directions, respectively, *p*_0_ is the constant term representing the mean qMRI, and *x*_0_, *y*_0_ and *z*_0_ are the Cartesian coordinates of the centre of mass of the ROI mask. This analysis was limited to the putamen and caudate nucleus as there were too few voxels for other structures to have an accurate estimation of the spatial gradient and because there have been reported spatial gradients for these regions in the past (Drori et al., 2022).

#### 2.2.4. Normative modelling

General Linear Model (GLM) analysis was conducted to investigate the impact of ageing and other demographic factors on qMRI parameters in various structures (ROI). The design matrix comprised four regressors: Age, Age^2^, Sex, and (left-/right-)Hemisphere:

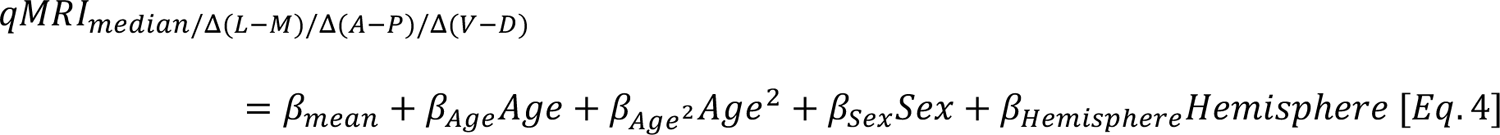

The utility of Age^2^ as a regressor is to consider the inverted-U shape effect observed in our R_1_ data which was also shown previously (Yeatman et al., 2014). Z-statistics for each regressor were computed by converting the t-statistic of GLM using *fsl_glm* (Jenkinson et al., 2012). In the following normative modelling analysis, the data were corrected for the sex-specific and inter-hemispherical effects based on the GLM results. Gaussian Process Regression (GRP) (Marquand et al., 2016b), a non-parametric supervised learning method, from PCNToolkit (Rutherford et al., 2022b) was used to compute the normative trajectory of the sex- and hemispheric-corrected qMRI metrics as a function of age with 10-fold cross-validation. To apply the normative model to the POM cohort, both R_2_* and *X* in the POM datasets were first corrected for the sex and hemispheric differences based on the GLM results of the ABRIM cohort, from which the z-statistics were computed using the GPR normative models. Two-sample t-tests were also performed to compare the z-scores of each structure between the healthy controls and Parkinson’s disease patients as a confirmatory analysis of previous findings where significantly higher values were found in substantia nigra (both R_2_* and *X*) and in globus pallidus (only *χ*) (Langkammer et al., 2016; Pyatigorskaya et al., 2020). The Benjamini and Hochberg procedure was used to control the false discovery rate (FDR) of 0.05 (Benjamini and Hochberg, 1995; Benjamini and Yekutieli, 2001). An adjusted *p*-value of 0.05 after the FDR correction is considered statistically significant. Furthermore, we compute the normative trajectories using only the POM healthy individual data following the same processing steps to evaluate if the normative trajectories were comparable when data with a narrower age range was used.

#### 2.2.5. Correlation analysis

While both iron deposition and myelin concentration in the brain can contribute to all the qMRI metrics, the qMRI metrics are known to have different sensitivities to the changes in the iron and myelin concentrations. To investigate if the various qMRI metrics possess similar/redundant information, Pearson’s correlation analysis was performed on the qMRI metric z-scores derived from the GRP of all subjects for each subcortical grey matter structure. In this way, we can investigate if the deviations from the norm in one qMRI metric are predictive of the deviations in another metric. Similarly, the associations among the subcortical grey matter structures on each qMRI metric were studied by performing Pearson’s correlation analysis on the GRP’s z-scores between the structures. With this analysis, we aimed to evaluate if deviations from the norm are independent across ROIs, or if there are any networks of nuclei where the relaxation values tend to jointly fluctuate.

## 3. Results

Figure 2 presents the z-statistics results of the GLM analysis conducted on the qMRI metrics for each subcortical ROI. Ageing (as represented by either Age of Age^2^ regressor) exhibits the most statistically significant effects on the measured qMRI metrics within the subcortical and thalamic nuclei (Figure 2a&b). Across all structures, a z-score of at least 3.86 is observed in at least one of the qMRI values on ageing. Notably, among the subcortical nuclei, the nucleus accumbens (NAcc) displays the weakest age-dependence on R_1_ and R_2_*, while the internal and external globus pallidus (GPi) and (GPe), despite being highly paramagnetic, exhibit the weakest age-dependence on *χ*. Conversely, red nucleus (RN) shows a strong age dependence across all three metrics. Generally, ageing manifests a positive linear effect (depicted in red, Figure 2a) and a negative quadratic effect (depicted in blue, Figure 2b) on the qMRI metrics for the subcortical nuclei. The Age^2^ effect is more pronounced on R_1_ and to a lesser extent on R_2_*. Moreover, among the three qMRI metrics, Age exerts a stronger impact on R_1_ compared to R_2_* and *χ*. Sex demonstrates the weakest effect on qMRI among the four demographic factors (Figure 2c), while interhemispheric differences exhibit notable effects on various structures (Figure 2d). Interhemispheric effects are particularly prominent in the NAcc and thalamic sub-nuclei (also thalamus as a whole), with interhemispheric differences exclusive to R_1_ observed in the caudate nucleus (Cau), substantia nigra part compacta (SNc), and ventral pallidum (VP). When focusing on thalamic sub-nuclei, similar age-dependence trends are observable for R_1_ and R_2_* (positive and negative z-scores for Age and Age^2^, respectively), whereas for *χ*, the linear age dependence displays varying polarities across nuclei.

**Figure 2:**
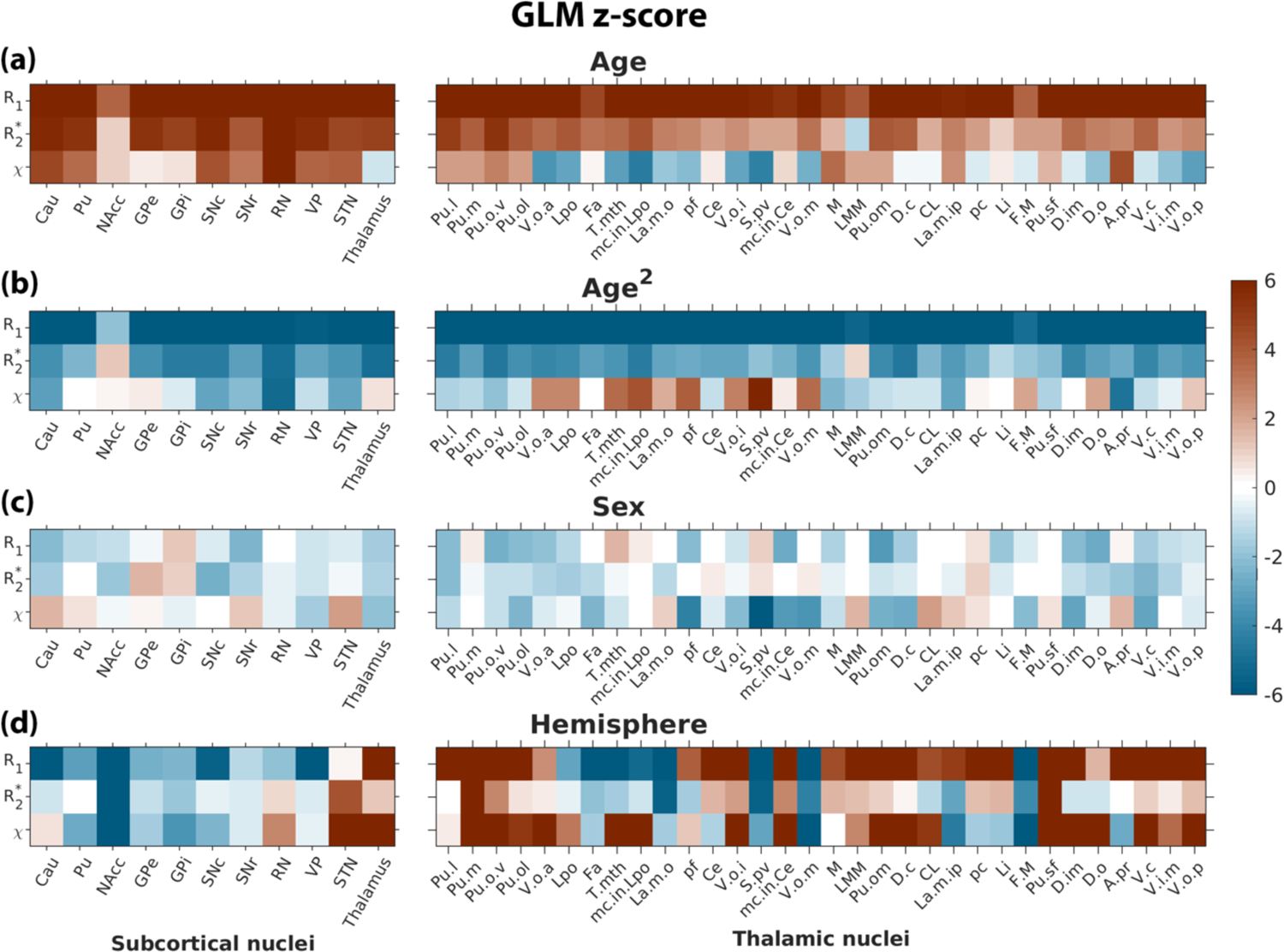
z-scores of GLM expressed in Eq. (3) for (a) Age, (b) Age^2^, (c) Sex and (d) Hemisphere. Within each panel, the z-scores for the subcortical nuclei (left) and thalamic sub-nuclei (right) are shown.

The normative trajectories on how subcortical nucleus volume, R_1_, R_2_* and *χ* change as a function of age are shown in Figure 3. Normalised tissue volumes generally decrease with increasing age, except for VP. The majority of R_1_ trajectories exhibit an inverted U-shape, whereas R_2_* and *χ* trajectories tend to be more linear, consistent with the GLM findings in Figure 2 despite the non-parametric nature of the GPR analysis. Notably, NAcc and VP stand out as clear outliers from the inverted U-shape trajectory of R_1_, displaying a parabolic behaviour that does not reach a maximum within the age range of our study. Conversely, most other nuclei appear to reach a maximum R_1_ between the ages of 45 and 60 years. Despite the flexibility of GPR, the derived trajectories support the utilisation of the quadratic age model within the age range here (18-79 years). Most structures exhibit both different medians and rates of development across the lifespan. Overall, spatially proximate sub-structures, such as GPe and GPi, and SNc and SNr, demonstrate similar behaviour both in volume and in R_1_, while the substantia nigra sub-structures are clearly distinct in both *χ* and R_2_* trajectories.

**Figure 3:**
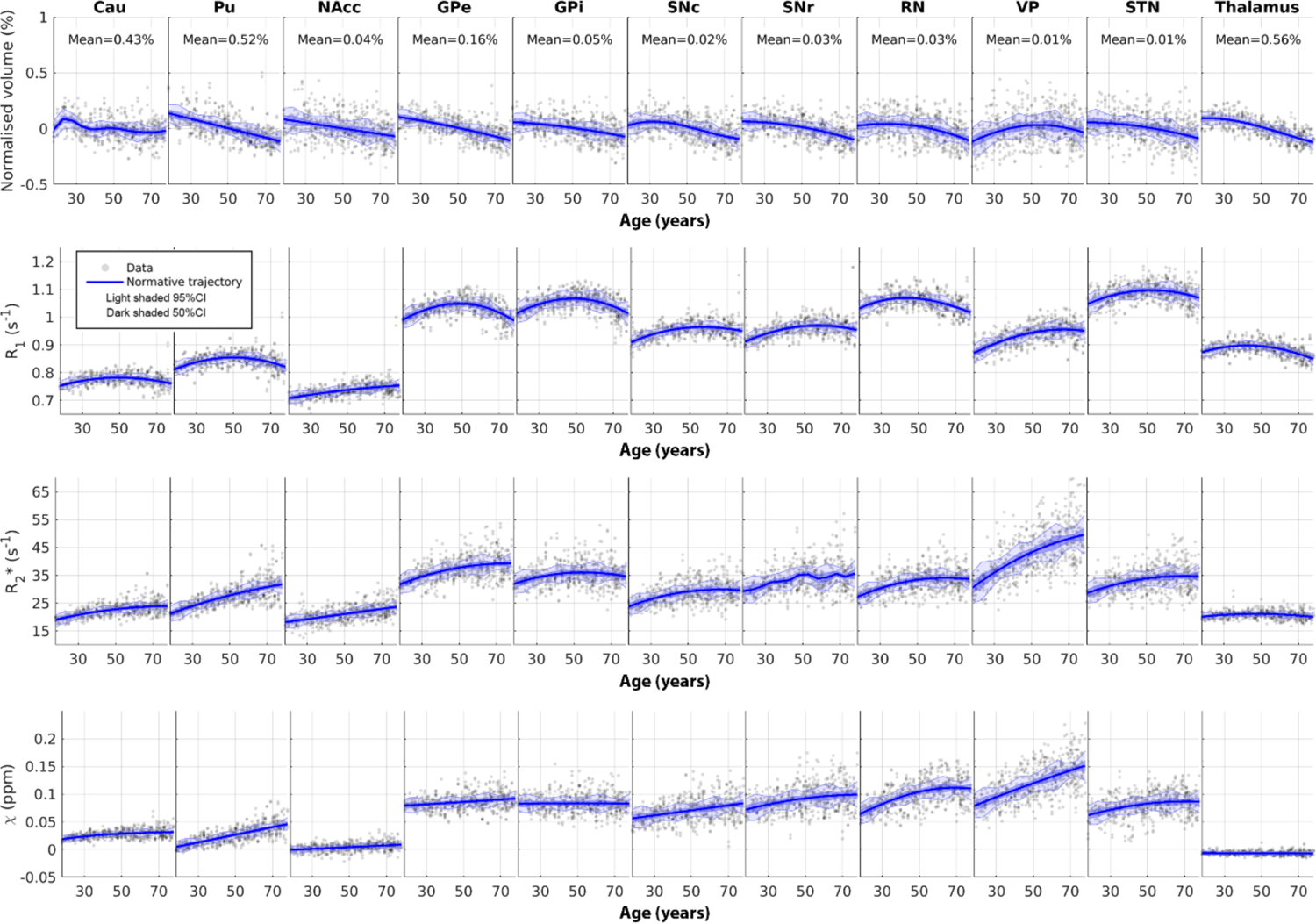
Normative models of normalised volume, R1, R2*, *χ* as a function of age for 11 different deep grey matter structures present in the MuSus-100 atlas. For visualisation purposes, the normalised volume of a given structure, *V*_’(),%_, is expressed as a percentage change of tissue volume in Eq. (2) and is shown at the top of each plot. The data points shown on the scatter plot are corrected for sex and hemispheric effects.

The trajectory of *χ* is nearly constant when treating the thalamus as a single structure (bottom right, Figure 3). However, further investigation of each thalamic sub-nucleus reveals that the nuclei have distinct trajectories (see Figure 4), and ageing shows opposite effects on *χ* in various structures. For example, the *χ* value increases as a function of age for nuclei close to the pulvinar region (bottom right, Figure 4), while an opposite trend is observed in the ventrallatero dorsal, ventral-anterior, and ventral-latero-ventral ROIs (top row, Figure 4). In contrast to *X*, R_1_ and R_2_* are relatively comparable among the thalamic sub-nuclei although they clearly have different means across the cohort population, suggesting that an alternative clustering of thalamic sub-nuclei could be obtained using mean R_1_ and R_2_* values. Among the thalamic sub-nuclei, the V.o.a (first column, top left, Figure 4) and the Pu.I (first column, bottom right, Figure 4) have the largest R_1_ and R_2_*, respectively, and the S.pv (fourth column, bottom left, Figure 4) and the Fa (third column, top left, Figure 4) have the lowest R_1_ and R_2_* values, respectively.

**Figure 4:**
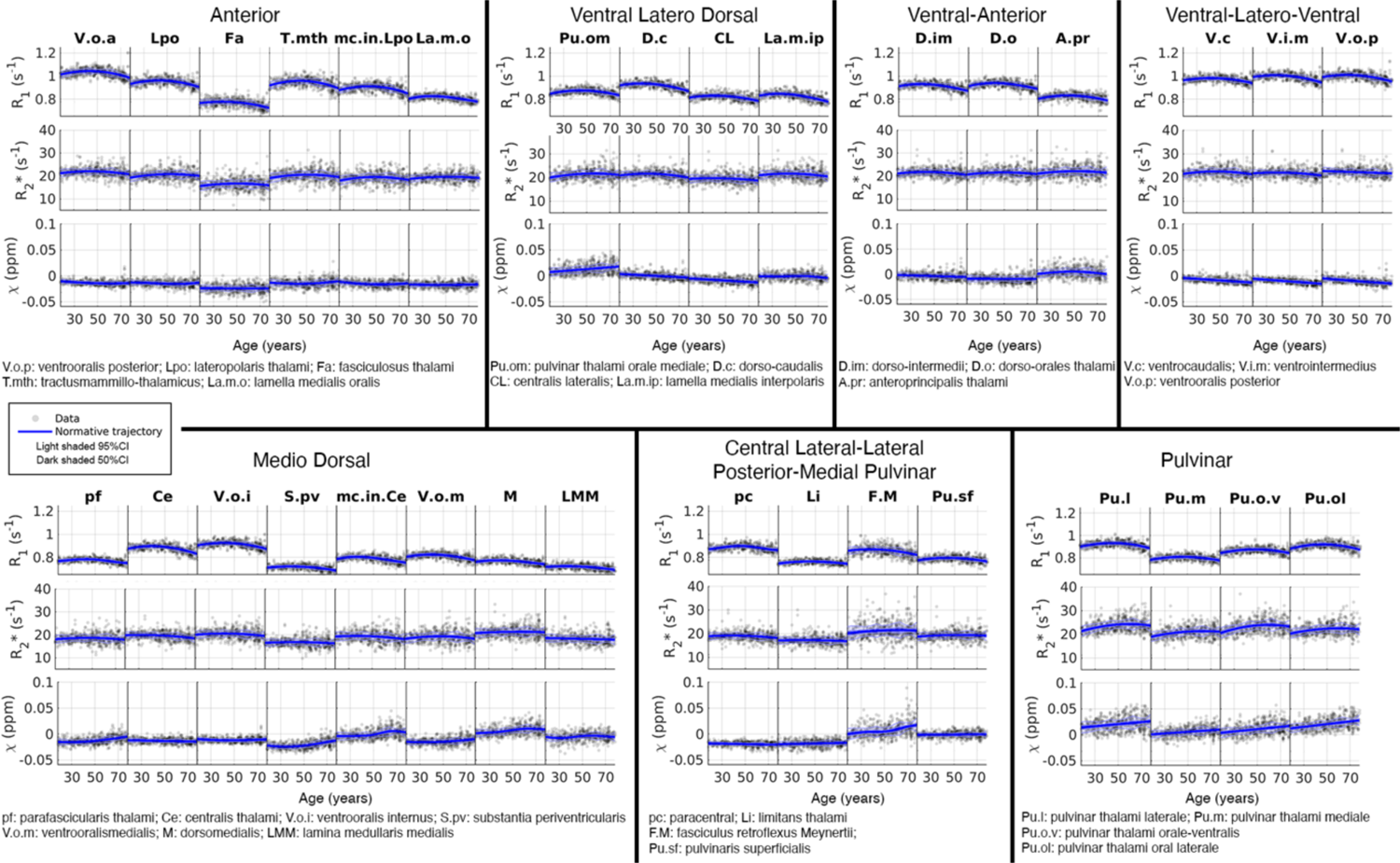
Normative models for the thalamic sub-nuclei defined in the MuSus-100 atlas as a function of age. To facilitate visualisation of the trajectories we have clustered these trajectories based on their anatomical proximity to the 7 thalamic subparts defined by the maximum likelihood atlas (Najdenovska et al., 2018).

The normative trajectories in Figures 3 and 4 provide an overview of the changes in qMRI metrics across adulthood for each nucleus. To delve deeper into how ageing may influence the spatial distribution of qMRI values and to reproduce previous findings (Drori et al., 2022), we extended the normative modelling analysis to the first-order spatial variation of the putamen and caudate nucleus in the MNI space. Figure 5a and 5b showcase the spatial variations on the group-averaged maps observed in the two structures, revealing a predominantly linear gradient. In the putamen, R_1_ exhibits a non-zero gradient offset in the Posterior-Anterior (P-A) and Ventral-Dorsal (V-D) directions, indicating spatial gradients present across all ages (top row Figure 5c), and these are not associated with the presence of distinct nuclei as in the thalamus. The age-dependent effect on the spatial gradient is evident in the Lateral-Medial (L-M) direction for R_2_* and *χ*, with the magnitude of the gradient increasing from medial to lateral with age (first column of Figure 5c). Similarly, age-dependent effects are observed in the P-A and V-D directions for R_2_*, albeit weaker compared to the L-M direction. In the caudate nucleus, a non-zero gradient offset is identified in the L-M direction for R_1_, with a parabolic change in gradient magnitude as a function of the subjects’ age (top corner, Figure 5d).

**Figure 5:**
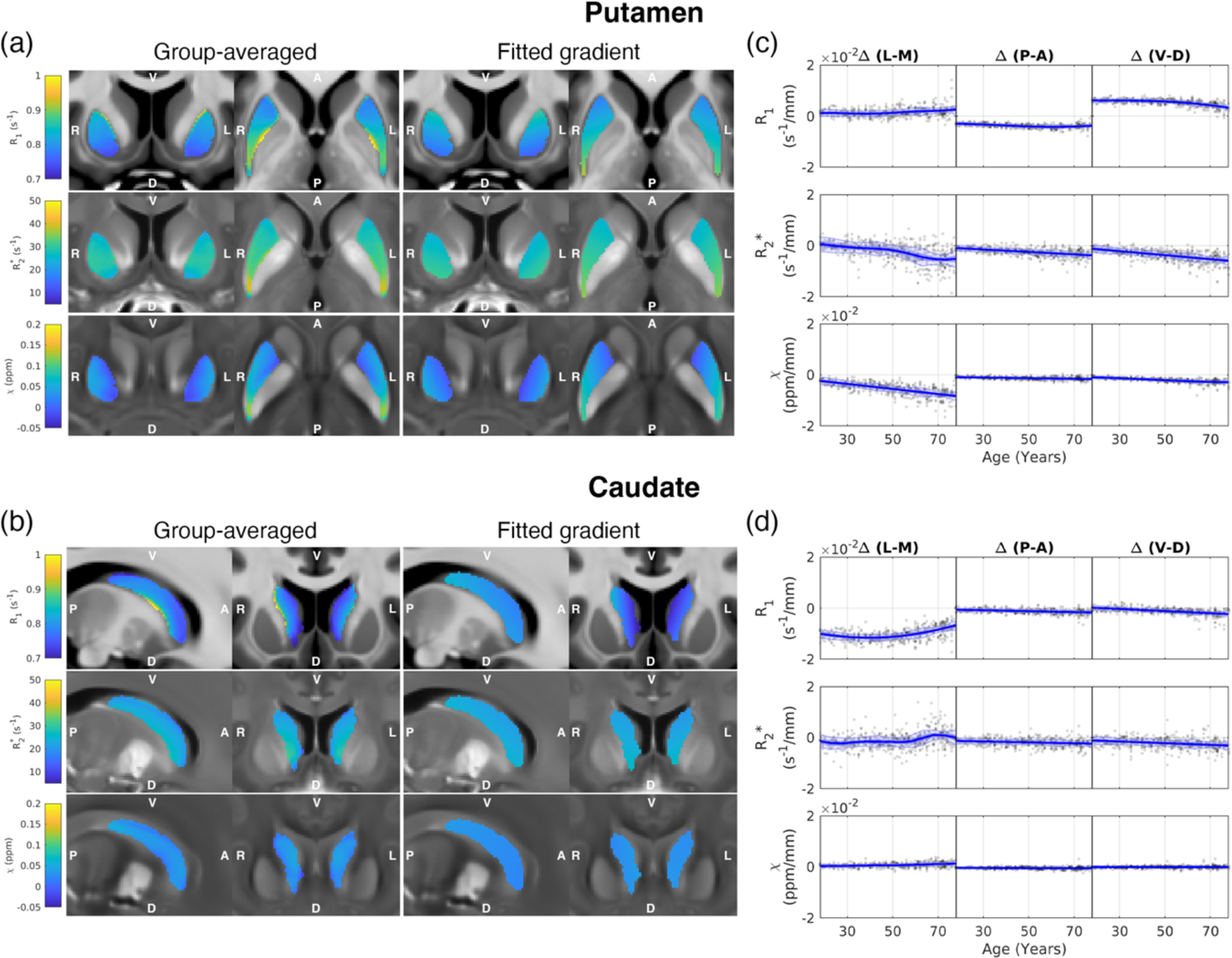
Example gradient overlaid on anatomical R1, R2* and *X* images in the (a) putamen and (b) caudate nucleus using group-averaged qMRI maps. The normative trajectories of the qMRI metrics in L-M, P-A, and V-D directions in the (c) putamen and (d) caudate nucleus.

Pearson’s correlation analysis of the z-scores between various qMRI metrics provides insights into their interrelationships. Weak-to-Moderate correlations (0.17≥R≥0.39) are observed between R_1_ and R_2_*, as well as between R_1_ and *χ* (0.10≥R≥0.34) across all subcortical nuclei, suggesting potential interdependencies in the mechanisms underlying these contrasts (Figure 6a). Moderate-to-strong correlations (0.44≥R≥0.81) between R_2_* and *χ* are observed in all nuclei except VP, indicating a linear relationship between changes in R_2_* and *χ* (third row, Figure 6a), after accounting for the large effects of age. While variations in R_2_* and *X* in subcortical nuclei are primarily driven by changes in iron concentration, single-compartment R_1_ is predominantly associated with MT effects, with some contributions from variations in iron concentration (Rooney et al., 2007). Conversely, changes in subcortical nucleus volume exhibit only weak correlations with their corresponding qMRI metrics (Figure 6a).

**Figure 6:**
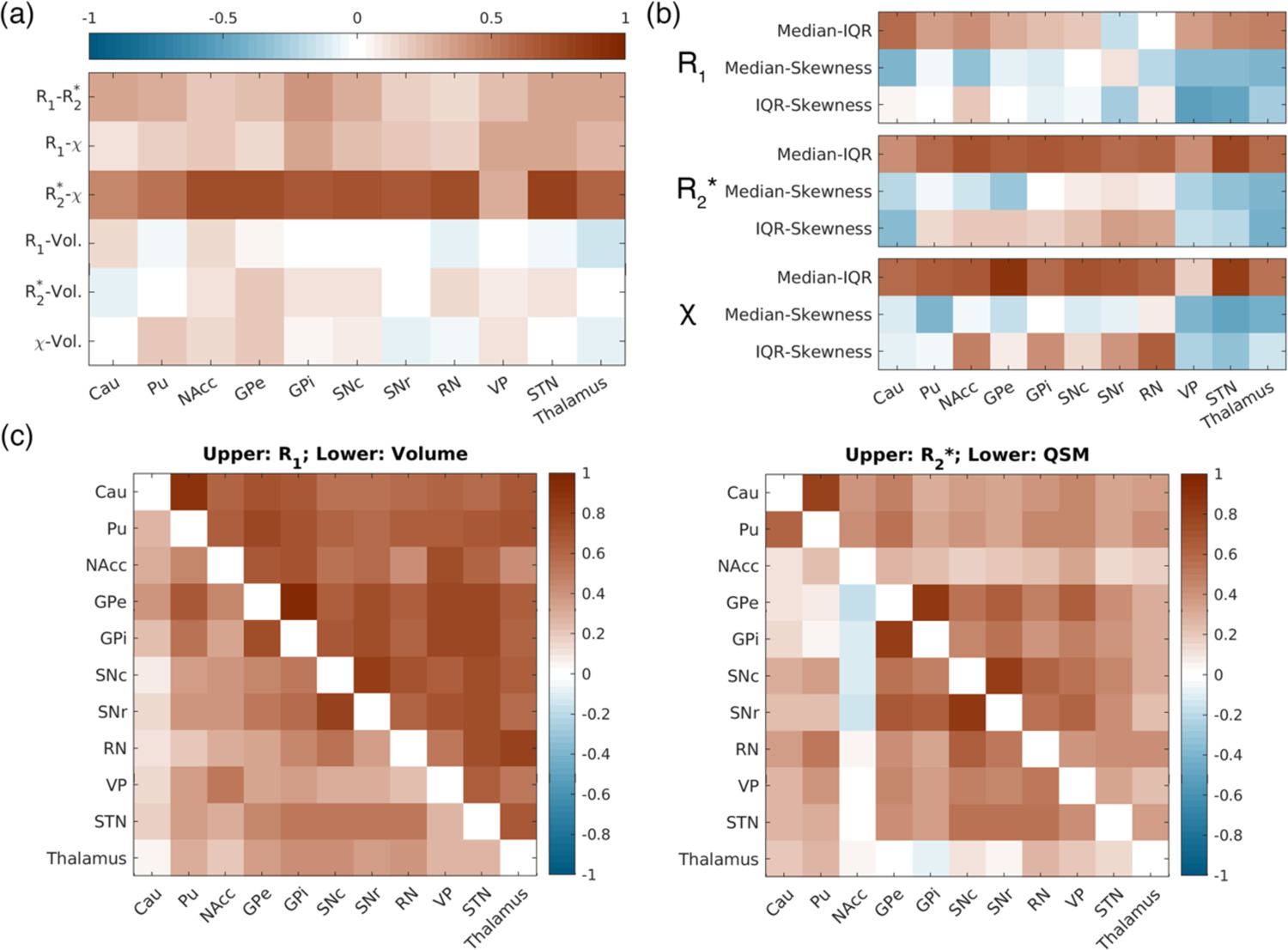
Shows various correlations between the z-scores derived from the normative models of both the volumetric and qMRI metrics: (a) Correlation between the z-scores across the four different metrics derived from Figure 4 (volume, R1, R2* and *X*) per ROI; b) Correlations within each qMRI metric between the z-scores of all subjects between the 3 statistical measures (median, IQR and skewness); c) and d) correlation matrices for volumetric and qMRI z-values across ROIs;

We observed that robust normative trajectories can also be extracted from the IQR of each nucleus (Figure S3), while the nuclei exhibit different offsets in skewness and remain relatively invariant across age with relatively strong variances (Figure S4). When examining the correlation between different univariate analysis metrics (such as median, IQR and skewness) of the same ROI and qMRI metric, Figure 6b reveals a notable correlation between median values and IQR (first row), with the correlation values comparable to those found between R_2_* and *X* in Figure 6a. Skewness demonstrates a reduced correlation with median and IQR, suggesting that skewness is dominated by noise variance rather than containing independent information when interpreted together with the findings in Figures S3 and S4.

Moving to the investigation of qMRI covariance networks, upon closer examination of individual metrics, R_1_ displays the strongest correlations in changes among the nuclei compared to other metrics (upper triangle, left panel, Figure 6c). Particularly strong correlations are observed in R_1_ between the caudate nucleus and putamen (R=0.87), GPe and GPi (R=0.95), and SNc and SNr (R=0.84). The strong correlations among all nuclei in R_1_ could be interpreted in two ways: (1) as a systematic bias per subject associated with the acquisition (e.g. increased or decreased flip angles not fully accounted for in the R_1_ quantification process); or (2) as an independent contrast mechanism that varies uniformly across all regions, e.g., increased macromolecular concentration or reduced hydration. The first hypothesis has been mitigated by factoring in the transmit field measurement in the computation of R_1_. In R_2_* and *χ*, two clusters of strong correlation are observed: one between the caudate nucleus and putamen (R=0.81 for R_2_* and R=0.61 for *χ*), and another among the globus pallidus, substantia nigra, red nucleus, and subthalamic nucleus (right panel, Figure 6c). This suggests that there might be two mechanistically different processes causing iron increases in the brain. The correlation matrix on volume changes is overall less similar to other metrics (lower triangle, left panel, Figure 6c), despite exhibiting the strongest correlation between GPe and GPi, and between SNc and SNr.

Finally, we explored the applicability of the R_2_* and *χ* normative trajectories using another dataset comprising both healthy individuals and Parkinson’s disease patients, characterised by a narrower age range in this cohort. When plotting the healthy individual data together with the normative models shown in Figure 3, most of the data points visually align with the normative trajectories (Figure 7a). Yet, the normative models derived from this narrower range of data (orange lines, Figure 7a) are less smooth when compared to having the wider age range data (blue lines, Figure 7a), particularly for the putamen, VP, and STN, suggesting the models may be overfitted and not account for the true age dependence. The robust fits of our normative models to the new data are highlighted by the distribution of the z-scores for all ROIs. The median z-scores are close to zero for almost all subcortical nuclei, and the z-score distributions of the healthy controls fall within the range of |z|<=1.96 (∼95% CI) for both R_2_* (yellow boxes, Figure 7b) and *χ* (yellow boxes, Figure 7c), though relatively stronger deviations are observed on R_2_* of the caudate nucleus, and *χ* of GPe, SNr, and RN. When examining early Parkinson’s patients, it is notable that most distributions remain centred around zeros, with exceptions noted for GPi and SNc – regions that are known to be associated with Parkinson’s disease (green boxes, Figure 7b&c). Two-sample t-test results indicate the z-scores of both R_2_* and *χ* in GPi are significantly higher in the Parkinson’s disease patients than in the healthy controls (adjusted *p*-values=6.8×10^-7^ and 0.0015 for R_2_* and *X*, respectively) and in SNc (adjusted *p*-values= 0.0058 and 0.018 for R_2_* and *X*, respectively).

**Figure 7:**
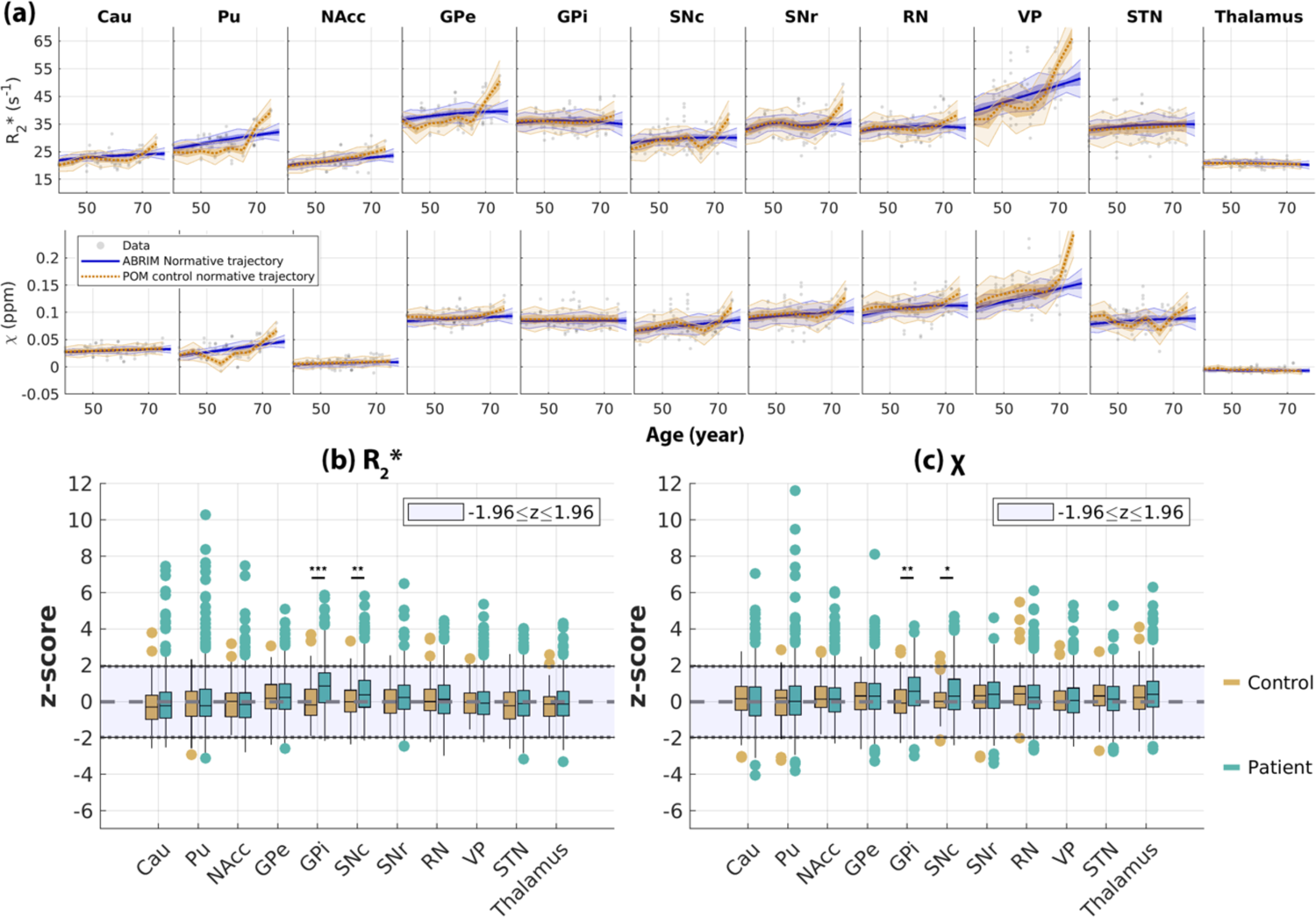
(a) Scatter plots of R2* (top row) and *X* (bottom row) values as a function of age in an independent sample of healthy controls (N=44) obtained from (Bloem et al., 2019). Blue and orange lines represent the normative models derived from the ABRIM study (as shown in Figure 3) and derived from the new control data. Panels (b) and (c) show the derived z-scores on the POM data using the ABRIM normative models for R2* and *X*, respectively. Within each panel, whisker plots are used to characterize the distribution (25, 50 and 75 percentile) of the controls (orange) and early Parkinson patients (green). \**p*-valueσ0.05; \*\**p*-valueσ0.01; \*\*\**p*-valueσ0.001.

## 5. Discussion

In this work, we demonstrate the effectiveness of integrating normative modelling with qMRI techniques at 3T. We believe that this study represents the most comprehensive single-study investigation to date of the variations in the three quantitative metrics (R_1_, R_2_* and *χ*) in the basal ganglia throughout adulthood. Our study benefits from a fairly large sample size of 260 healthy participants, with a balanced distribution of age and sex. This has allowed us to explore not only the evolution of these metrics with age beyond the simple young versus old adult comparison but also their heteroscedasticity, i.e., how the variance changes across different age groups. Prior studies focusing on R_1_, R_2_* and *χ* have primarily utilised 7T data and typically involved smaller sample sizes (Keuken et al., 2017). While larger studies at 3T do exist, they have mainly focused on R_2_* and *χ* (Li et al., 2021; Treit et al., 2021), or covered a broader age range (Li et al., 2014), including paediatric populations. The age range below 10 years is often characterised by rapid changes, making it challenging to observe and model the subtler variations that occur in adulthood. Our study addresses this gap by focusing on the adult population.

The inverted U-shaped trajectory observed in R_1_ has been in the past linked to processes of myelination and demyelination when investigating white matter (Yeatman et al., 2014). While refraining from overinterpreting the significance of R_1_ and its association with macromolecular content or myelination, it is noteworthy that the peak age in our normative trajectories typically falls between 50 and 65 years, considerably later than the observations in white matter bundles (Yeatman et al., 2014). One explanation for this shift in the maturation peak measured in R_1_ could be attributed to the overlap of a myelination trajectory that peaks at an earlier age, with a linear increase of iron deposition. Such a shift would not be observed in R_1_ of white matter where iron concentrations are orders of magnitude lower. Similar age trends are generally observable in R_1_ across subcortical nuclei, and to a lesser extent in R_2_* (as evidenced by the differences in curvatures in R_1_ and R_2_* in Figure 3), while *χ* appears to be less influenced by the myelination process in subcortical nuclei. It is important to note that *χ* has been identified as highly sensitive to myelination in white matter (Liu et al., 2011; Lodygensky et al., 2012). This sensitivity is primarily attributed to microstructural effects that are amplified in tightly aligned white matter bundles (Luo et al., 2014; Wharton and Bowtell, 2015), which are less prevalent in subcortical nuclei.

It is well-established that R_2_* and *χ* encode similar contrast mechanisms albeit in different ways (Stüber et al., 2014), and they are particularly sensitive to iron concentration when examining subcortical nuclei (Langkammer et al., 2012). This similarly is underscored by the strong resemblance of the normative models depicted in Figure 3 (3^rd^ and 4^th^ rows). There is a discernible association between regions with higher and lower values and the rates at which they change throughout adulthood. Even after removing the age confound and transforming the metrics of interest to z-statistics, our correlation analysis depicted in Figure 6a indicates a high level of interrelation between these two parameters (mean correlation R_2_* vs *χ* = 0.64). Although the correlation of these values to the z-scores of R_1_ is diminished (mean correlations of R_1_ vs R_2_*=0.28; R_1_ vs *χ*=0.23), it confirms that iron contributes weakly to the contrast observed in subcortical nuclei. In Supplementary Figures S2-S4 and Figure 6b, we have demonstrated those statistical measures beyond the median and the spatial gradients, such as skewness. While these measures are relatively noisier than the median, they have potential to serve as additional imaging biomarkers when combining with local spatial information (Li et al., 2019). Volumetry is frequently utilised in ageing and cohort studies (Bethlehem et al., 2022; Rutherford et al., 2022a), yet our results suggest that the process of volume change is largely distinct from the microstructural alterations (including macromolecular concentration and iron deposition) measured by the three qMRI methods addressed herein. Future studies to explore the association between qMRI metrics and behavioural data available in the ABRIM data collection (Jansen et al., 2024) may provide valuable insight into cognitive ageing with (micro)structural alterations.

Our examination of the normative trajectories of various thalamic sub-nuclei, as depicted in Figure 4, yields an unexpected observation. We found that while the trajectories of R_2_* either remained constant or exhibited minor increases, the *χ* values showed small but noticeable increases or decreases with age, varying across different nuclei. One possible explanation for these changes, beyond the susceptibility changes resulting from the actual structural alteration, could be related to arbitrary susceptibility reference used in the QSM processing (QSM Consensus Organization Committee et al., 2024). If the mean of the absolute magnetic susceptibility of the whole brain would be increasing at a higher rate than that of those thalamic sub-nuclei (D.c, CL, V.c, V.i.m and V.o.p.), the fact we assume that the mean brain susceptibility of the brain remains equal to zero would result in an apparent decrease of the measured relative susceptibility.

Spatial variation in R_1_ and *χ* within the putamen and caudate nucleus have been previously documented by other research teams, with variation observed between young and elderly individuals in both R_1_ and *χ* (Keuken et al., 2017). This spatial gradient may hold significance in the context of Parkinson’s disease (Keuken et al., 2017), as alterations in the anterior-posterior (A-P) gradient in R_1_ maps have been observed in Parkinson’s patients compared to age-matched healthy individuals (Drori et al., 2022). Furthermore, the tails of the Putamen in *X* maps have been found to be significantly associated with decreased motor performance in Parkinson’s patients (Drori et al., 2022).

Here, we opted to conduct normative modelling using high SNR approaches derived from atlas-based parcellation. The atlas used here (MuSus-100) was constructed using similar qMRI maps. To optimise the coregistration between our dataset and the associated templates from the atlases, we implemented a robust method to address the image contrast variations between site-specific data and the atlas template. We also extended our analysis to another atlas (Alkemade et al., 2020) to demonstrate the robustness of this approach for subcortical parcellation (see Supplementary Figure S2). The contrast variations in the hybrid image may arise due to differences in MRI scanner field strength, imaging sequences, or the signal model used to derive the qMRI values, potentially introducing image intensity biases between datasets. To streamline the parcellation process based on volumetric image registration, we integrated Step 1 of the co-registration procedure into the SEPIA toolbox (Chan and Marques, 2021), aiming to facilitate the widespread adoption of this methodology.

While our focus has been on examining normative models for individual ROIs, it is essential to acknowledge that this information is not entirely independent across the brain. We observed correlation values exceeding 0.5 for almost all ROIs for R_1_. This could either indicate a potential consistent measurement bias, such as inaccurate B_1_ compensation during R_1_ computation or suggest that there is a common macromolecular change influencing all brain regions similarly due to co-development or co-regulation (Carmon et al., 2020). Despite our careful consideration of the B_1_ effect (Marques and Gruetter, 2013), there may still be residual biases due to differences in inversion efficiency among subjects, where the system might have underestimated or overestimated the reference voltage. For the R_2_* and *χ* z-scores, strong correlations were observed between the caudate nucleus and putamen, as well as among the sub-structures of the globus pallidus and substantia nigra (R>0.6), possibly driven by their close spatial proximity. By reducing the correlation threshold to 0.4, we can visually discern two main “networks of variations”: one comprising the caudate nucleus and putamen, and the other including the globus pallidus, substantia nigra, red nucleus, ventral pallidum, and subthalamic nucleus.

We emphasise the significance of creating normative models for each quantitative metric and ROI to understand these metrics within the context of a population, it is also important to recognise that these derived models, even within the realm of quantitative imaging, may not hold universal validity across different studies. Variations in acquisition parameters, including repetition times, echo times, inversion times, image resolution, and vendor hardware, as well as differences in the exact pipeline implementation to derive ROI metrics, are bound to introduce study biases (Karakuzu et al., 2022; Teixeira et al., 2019). Therefore, when combining datasets from different sites, it is crucial to consider harmonisation procedures akin to those commonly employed in other morphometric analyses (Bayer et al., 2022; Fortin et al., 2017; Kia et al., 2020; Pomponio et al., 2020). Both the raw quantitative maps and the derived quantitative metrics from our healthy volunteers, encompassing various subcortical nuclei and atlases evaluated here, have been made publicly available to facilitate integration into subsequent research endeavours (Jansen et al., 2024).

## 6. Conclusion

In this study, we conducted a comprehensive normative modelling analysis of R_1_, R_2_*, and *X* in subcortical nuclei and thalamic sub-nuclei across adulthood. Our analysis utilised a cohort with an evenly distributed age and sex ratio. The normative models we derived take into account the changes in variance across age, thereby enabling the derivation of z-statistics with new data. Interestingly, the shapes of the trajectories align with existing literature findings, despite our use of a non-parametric approach to compute these models. The observed differences in the trajectory shapes between R_1_ and R_2_*/*χ* suggest that the mechanisms driving these contrasts are not identical. We further confirmed that the putamen and caudate nucleus exhibit an age-dependent spatial distribution of these qMRI metrics. This finding could potentially pave the way for more in-depth studies of spatial distribution changes in neurological disorders, such as Parkinson’s disease.

## Abbreviations

Pf: Parafascicularis thalami

pc: Paracentral

Ce: Centralis thalami

Pu.l: Pulvinar thalami laterale

Pu.m: Pulvinar thalami mediale

Pu.o.v: Pulvinar thalami orale-ventralis

V.c: Ventrocaudalis

Li: Limitans thalami

V.i.m: Ventrointermedius

V.o.p: Ventrooralis posterior

V.o.a: Ventrooralis anterior

V.o.i: Ventrooralis internus

Lpo: Lateropolaris thalami

Fa: Fasciculosus thalami

T.mth: Tractusmammillo-thalamicus mc in Lpoc

La.m.o: Lamella medialis oralis

S.pv: Substantia periventricularis mc in Ce

F.M: Fasciculus retroflexus Meynertii

V.o.m: Ventrooralismedialis

Pu.ol: Pulvinar thalami oral laterale

Pu.om: Pulvinar thalami orale mediale

Pu.sf: Pulvinaris superficialis

D.c: Dorso-caudalis

D.im: Dorso-intermedii

D.o: Dorso-orales thalami

M: Dorsomedialis

A.pr: Anteroprincipalis thalami

CL: Centralis lateralis

La.m.ip: Lamella medialis interpolaris

LMM: Lamina medullaris medialis

Cau: Caudate nucleus

Pu: Putamen

GPe: External globus pallidus

GPi: Internal globus pallidus

NAcc: Nucleus accumbens

SNr: Substantia nigra pars reticulata

SNc: Substantia nigra pars compacta

RN: Red nucleus

VP: Ventral pallidum

STN: Subthalamic nucleus

SD: Standard deviation

TI: Inversion time

TR: Repetition time

TE: Echo time

TA: Acquisition time

GRE: Gradient echo imaging

CI: Confidence interval

## Data and Code Availability

The data and code that support the findings of this study will be made available upon the acceptance of the manuscript.

## Author Contributions

Kwok-Shing Chan: Conceptualization, Methodology, Software, Validation, Formal analysis, Investigation, Writing – Original Draft, Visualization

Marcel P. Zwiers: Software, Resources, Data Curation, Writing – Review & Editing

Michelle G. Jansen: Data Curation, Writing – Review & Editing

Joukje M. Oosterman: Project administration, Funding acquisition, Writing – Review & Editing

David G. Norris: Supervision, Project administration, Funding acquisition, Writing – Review & Editing

Christian F. Beckmann: Conceptualization, Supervision, Funding acquisition

José P. Marques: Conceptualization, Methodology, Supervision, Writing – Original Draft, Visualization

## Acknowledgements

The ABRIM study was funded by the European FP7 program, FP7-PEOPLE-2013-ITN, Marie-Curie Action, “Initial Training Networks” named “Advanced Brain Imaging with MRI” (no. 608123). CFB acknowledges funding from the Wellcome Trust Collaborative Award in Science 215573/Z/19/Z and the Netherlands Organization for Scientific Research Vici Grant No. 17854. The authors thank Dr. Rick Helmich and Dr. Martin Johansson for the access to the POM datasets.

## References

Acosta-Cabronero, J., Betts, M.J., Cardenas-Blanco, A., Yang, S., Nestor, P.J., 2016. In Vivo MRI Mapping of Brain Iron Deposition across the Adult Lifespan. J Neurosci 36, 364–374. 10.1523/jneurosci.1907-15.2016

Acosta-Cabronero, J., Williams, G.B., Cardenas-Blanco, A., Arnold, R.J., Lupson, V., Nestor, P.J., 2013. In Vivo Quantitative Susceptibility Mapping (QSM) in Alzheimer’s Disease. Plos One 8, e81093. 10.1371/journal.pone.0081093

Alkemade, A., Mulder, M.J., Groot, J.M., Isaacs, B.R., van Berendonk, N., Lute, N., Isherwood, S.J., Bazin, P.-L., Forstmann, B.U., 2020. The Amsterdam Ultra-high field adult lifespan database (AHEAD): A freely available multimodal 7 Tesla submillimeter magnetic resonance imaging database. Neuroimage 221, 117200. 10.1016/j.neuroimage.2020.117200

Avants, B.B., Tustison, N.J., Song, G., Cook, P.A., Klein, A., Gee, J.C., 2011. A reproducible evaluation of ANTs similarity metric performance in brain image registration. Neuroimage 54, 2033–2044. 10.1016/j.neuroimage.2010.09.025

Bayer, J.M.M., Thompson, P.M., Ching, C.R.K., Liu, M., Chen, A., Panzenhagen, A.C., Jahanshad, N., Marquand, A., Schmaal, L., Sämann, P.G., 2022. Site effects how-to and when: An overview of retrospective techniques to accommodate site effects in multi-site neuroimaging analyses. Front. Neurol. 13, 923988. 10.3389/fneur.2022.923988

Benjamini, Y., Hochberg, Y., 1995. Controlling the False Discovery Rate: A Practical and Powerful Approach to Multiple Testing. J. R. Stat. Soc.: Ser. B (Methodol.) 57, 289–300. 10.1111/j.2517-6161.1995.tb02031.x

Benjamini, Y., Yekutieli, D., 2001. The control of the false discovery rate in multiple testing under dependency. Ann. Stat. 29. 10.1214/aos/1013699998

Bethlehem, R.A.I., Seidlitz, J., White, S.R., Vogel, J.W., Anderson, K.M., Adamson, C., Adler, S., Alexopoulos, G.S., Anagnostou, E., Areces-Gonzalez, A., Astle, D.E., Auyeung, B., Ayub, M., Bae, J., Ball, G., Baron-Cohen, S., Beare, R., Bedford, S.A., Benegal, V., Beyer, F., Blangero, J., Cábez, M.B., Boardman, J.P., Borzage, M., Bosch-Bayard, J.F., Bourke, N., Calhoun, V.D., Chakravarty, M.M., Chen, C., Chertavian, C., Chetelat, G., Chong, Y.S., Cole, J.H., Corvin, A., Costantino, M., Courchesne, E., Crivello, F., Cropley, V.L., Crosbie, J., Crossley, N., Delarue, M., Delorme, R., Desrivieres, S., Devenyi, G.A., Biase, M.A.D., Dolan, R., Donald, K.A., Donohoe, G., Dunlop, K., Edwards, A.D., Elison, J.T., Ellis, C.T., Elman, J.A., Eyler, L., Fair, D.A., Feczko, E., Fletcher, P.C., Fonagy, P., Franz, C.E., Galan-Garcia, L., Gholipour, A., Giedd, J., Gilmore, J.H., Glahn, D.C., Goodyer, I.M., Grant, P.E., Groenewold, N.A., Gunning, F.M., Gur, R.E., Gur, R.C., Hammill, C.F., Hansson, O., Hedden, T., Heinz, A., Henson, R.N., Heuer, K., Hoare, J., Holla, B., Holmes, A.J., Holt, R., Huang, H., Im, K., Ipser, J., Jack, C.R., Jackowski, A.P., Jia, T., Johnson, K.A., Jones, P.B., Jones, D.T., Kahn, R.S., Karlsson, H., Karlsson, L., Kawashima, R., Kelley, E.A., Kern, S., Kim, K.W., Kitzbichler, M.G., Kremen, W.S., Lalonde, F., Landeau, B., Lee, S., Lerch, J., Lewis, J.D., Li, J., Liao, W., Liston, C., Lombardo, M.V., Lv, J., Lynch, C., Mallard, T.T., Marcelis, M., Markello, R.D., Mathias, S.R., Mazoyer, B., McGuire, P., Meaney, M.J., Mechelli, A., Medic, N., Misic, B., Morgan, S.E., Mothersill, D., Nigg, J., Ong, M.Q.W., Ortinau, C., Ossenkoppele, R., Ouyang, M., Palaniyappan, L., Paly, L., Pan, P.M., Pantelis, C., Park, M.M., Paus, T., Pausova, Z., Paz-Linares, D., Binette, A.P., Pierce, K., Qian, X., Qiu, J., Qiu, A., Raznahan, A., Rittman, T., Rodrigue, A., Rollins, C.K., Romero-Garcia, R., Ronan, L., Rosenberg, M.D., Rowitch, D.H., Salum, G.A., Satterthwaite, T.D., Schaare, H.L., Schachar, R.J., Schultz, A.P., Schumann, G., Schöll, M., Sharp, D., Shinohara, R.T., Skoog, I., Smyser, C.D., Sperling, R.A., Stein, D.J., Stolicyn, A., Suckling, J., Sullivan, G., Taki, Y., Thyreau, B., Toro, R., Traut, N., Tsvetanov, K.A., Turk-Browne, N.B., Tuulari, J.J., Tzourio, C., Vachon-Presseau, É., Valdes-Sosa, M.J., Valdes-Sosa, P.A., Valk, S.L., van Amelsvoort, T., Vandekar, S.N., Vasung, L., Victoria, L.W., Villeneuve, S., Villringer, A., Vértes, P.E., Wagstyl, K., Wang, Y.S., Warfield, S.K., Warrier, V., Westman, E., Westwater, M.L., Whalley, H.C., Witte, A.V., Yang, N., Yeo, B., Yun, H., Zalesky, A., Zar, H.J., Zettergren, A., Zhou, J.H., Ziauddeen, H., Zugman, A., Zuo, X.N., 3R-BRAIN, Rowe, C., Initiative, A.D.N., Frisoni, G.B., Team, C., Cam-CAN, CCNP, COBRE, cVEDA, Group, E.D.B.A.W., Project, D.H.C., FinnBrain, Study, H.A.B., IMAGEN, KN E96, Aging, T.M.C.S. of, NSPN, POND, Binette, A.P., VETSA, Bullmore, E.T., Alexander-Bloch, A.F., 2022. Brain charts for the human lifespan. Nature 604, 525–533. 10.1038/s41586-022-04554y

Bloem, B.R., Marks, W.J., de Lima, A.L.S., Kuijf, M.L., van Laar, T., Jacobs, B.P.F., Verbeek, M.M., Helmich, R.C., van de Warrenburg, B.P., Evers, L.J.W., intHout, J., van de Zande, T., Snyder, T.M., Kapur, R., Meinders, M.J., 2019. The Personalized Parkinson Project: examining disease progression through broad biomarkers in early Parkinson’s disease. BMC Neurol. 19, 160. 10.1186/s12883-019-1394-3

Bookheimer, S.Y., Salat, D.H., Terpstra, M., Ances, B.M., Barch, D.M., Buckner, R.L., Burgess, G.C., Curtiss, S.W., Diaz-Santos, M., Elam, J.S., Fischl, B., Greve, D.N., Hagy, H.A., Harms, M.P., Hatch, O.M., Hedden, T., Hodge, C., Japardi, K.C., Kuhn, T.P., Ly, T.K., Smith, S.M., Somerville, L.H., Uğurbil, K., van der Kouwe, A., Essen, D.V., Woods, R.P., Yacoub, E., 2019. The Lifespan Human Connectome Project in Aging: An overview. NeuroImage 185, 335–348. 10.1016/j.neuroimage.2018.10.009

Burgetova, R., Dusek, P., Burgetova, A., Pudlac, A., Vaneckova, M., Horakova, D., Krasensky, J., Varga, Z., Lambert, L., 2021. Age-related magnetic susceptibility changes in deep grey matter and cerebral cortex of normal young and middle-aged adults depicted by whole brain analysis. Quant. Imaging Med. Surg. 0, 0–0. 10.21037/qims-21-87

Carmon, J., Heege, J., Necus, J.H., Owen, T.W., Pipa, G., Kaiser, M., Taylor, P.N., Wang, Y., 2020. Reliability and comparability of human brain structural covariance networks. NeuroImage 220, 117104. 10.1016/j.neuroimage.2020.117104

Chan, K.-S., Marques, J.P., 2021. SEPIA—Susceptibility mapping pipeline tool for phase images. Neuroimage 227, 117611. 10.1016/j.neuroimage.2020.117611

Costa, S.D., van der Zwaag, W., Marques, J.P., Frackowiak, R.S.J., Clarke, S., Saenz, M., 2011. Human Primary Auditory Cortex Follows the Shape of Heschl’s Gyrus. J. Neurosci. 31, 14067–14075. 10.1523/jneurosci.2000-11.2011

Deh, K., Nguyen, T.D., Eskreis-Winkler, S., Prince, M.R., Spincemaille, P., Gauthier, S., Kovanlikaya, I., Zhang, Y., Wang, Y., 2015. Reproducibility of quantitative susceptibility mapping in the brain at two field strengths from two vendors. J. Magn. Reson. Imaging 42, 1592–1600. 10.1002/jmri.24943

Drori, E., Berman, S., Mezer, A.A., 2022. Mapping microstructural gradients of the human striatum in normal aging and Parkinson’s disease. Sci Adv 8, eabm1971. 10.1126/sciadv.abm1971

Dymerska, B., Eckstein, K., Bachrata, B., Siow, B., Trattnig, S., Shmueli, K., Robinson, S.D., 2021. Phase unwrapping with a rapid opensource minimum spanning tree algorithm (ROMEO). Magn. Reson. Med. 85, 2294–2308. 10.1002/mrm.28563

Erramuzpe, A., Schurr, R., Yeatman, J.D., Gotlib, I.H., Sacchet, M.D., Travis, K.E., Feldman, H.M., Mezer, A.A., 2020. A Comparison of Quantitative R1 and Cortical Thickness in Identifying Age, Lifespan Dynamics, and Disease States of the Human Cortex. Cereb Cortex 31, 1211–1226. 10.1093/cercor/bhaa288

Fonov, V., Evans, A., McKinstry, R., Almli, C., Collins, D., 2009. Unbiased nonlinear average age-appropriate brain templates from birth to adulthood. NeuroImage 47, S102. 10.1016/s1053-8119(09)70884-5

Fortin, J.-P., Parker, D., Tunç, B., Watanabe, T., Elliott, M.A., Ruparel, K., Roalf, D.R., Satterthwaite, T.D., Gur, R.C., Gur, R.E., Schultz, R.T., Verma, R., Shinohara, R.T., 2017. Harmonization of multi-site diffusion tensor imaging data. NeuroImage 161, 149–170. 10.1016/j.neuroimage.2017.08.047

Ghadery, C., Pirpamer, L., Hofer, E., Langkammer, C., Petrovic, K., Loitfelder, M., Schwingenschuh, P., Seiler, S., Duering, M., Jouvent, E., Schmidt, H., Fazekas, F., Mangin, J.-F., Chabriat, H., Dichgans, M., Ropele, S., Schmidt, R., 2015. R2* mapping for brain iron: associations with cognition in normal aging. Neurobiol Aging 36, 925–932. 10.1016/j.neurobiolaging.2014.09.013

Gil, R., Khabipova, D., Zwiers, M., Hilbert, T., Kober, T., Marques, J.P., 2016. An in vivo study of the orientation-dependent and independent components of transverse relaxation rates in white matter. NMR in biomedicine 29, 1780–1790. 10.1002/nbm.3616

Gur, R.C., Gunning-Dixon, F.M., Turetsky, B.I., Bilker, W.B., Gur, R.E., 2002. Brain Region and Sex Differences in Age Association With Brain Volume: A Quantitative MRI Study of Healthy Young Adults. Am. J. Geriatr. Psychiatry 10, 72–80. 10.1097/00019442-200201000-00009

He, C., Guan, X., Zhang, W., Li, J., Liu, C., Wei, H., Xu, X., Zhang, Y., 2022. Quantitative susceptibility atlas construction in Montreal Neurological Institute space: towards histological-consistent iron-rich deep brain nucleus subregion identification. Brain Struct Funct 1–23. 10.1007/s00429-022-02547-1

Hoopes, A., Mora, J.S., Dalca, A.V., Fischl, B., Hoffmann, M., 2022. SynthStrip: skull-stripping for any brain image. NeuroImage 260, 119474. 10.1016/j.neuroimage.2022.119474

Jack, C.R., Bernstein, M.A., Fox, N.C., Thompson, P., Alexander, G., Harvey, D., Borowski, B., Britson, P.J., Whitwell, J.L., Ward, C., Dale, A.M., Felmlee, J.P., Gunter, J.L., Hill, D.L.G., Killiany, R., Schuff, N., Fox-Bosetti, S., Lin, C., Studholme, C., DeCarli, C.S., Krueger, G., Ward, H.A., Metzger, G.J., Scott, K.T., Mallozzi, R., Blezek, D., Levy, J., Debbins, J.P., Fleisher, A.S., Albert, M., Green, R., Bartzokis, G., Glover, G., Mugler, J., Weiner, M.W., 2008. The Alzheimer’s disease neuroimaging initiative (ADNI): MRI methods. J. Magn. Reson. Imaging 27, 685–691. 10.1002/jmri.21049

Jäncke, L., Mérillat, S., Liem, F., Hänggi, J., 2015. Brain size, sex, and the aging brain. Hum. Brain Mapp. 36, 150–169. 10.1002/hbm.22619

Jansen, M.G., Zwiers, M.P., Marques, J.P., Chan, K.-S., Amelink, J.S., Altgassen, M., Oosterman, J.M., Norris, D.G., 2024. The Advanced BRain Imaging on ageing and Memory (ABRIM) data collection: Study design, data processing, and rationale. PLOS ONE 19, e0306006. 10.1371/journal.pone.0306006

Jenkinson, M., Beckmann, C.F., Behrens, T.E.J., Woolrich, M.W., Smith, S.M., 2012. FSL. Neuroimage 62, 782–790. 10.1016/j.neuroimage.2011.09.015

Karakuzu, A., Biswas, L., Cohen-Adad, J., Stikov, N., 2022. Vendor-neutral sequences and fully transparent workflows improve inter-vendor reproducibility of quantitative MRI. Magn. Reson. Med. 88, 1212–1228. 10.1002/mrm.29292

Keuken, M.C., Bazin, P.-L., Backhouse, K., Beekhuizen, S., Himmer, L., Kandola, A., Lafeber, J.J., Prochazkova, L., Trutti, A., Schäfer, A., Turner, R., Forstmann, B.U., 2017. Effects of aging on, and QSM MRI values in the subcortex. Brain Struct Funct 222, 2487–2505. 10.1007/s00429-016-1352-4

Kia, S.M., Huijsdens, H., Dinga, R., Wolfers, T., Mennes, M., Andreassen, O.A., Westlye, L.T., Beckmann, C.F., Marquand, A.F., 2020. Hierarchical Bayesian Regression for Multi-site Normative Modeling of Neuroimaging Data. Springer International Publishing, pp. 699–709.

Kiersnowski, O.C., Karsa, A., Wastling, S.J., Thornton, J.S., Shmueli, K., 2023. Investigating the effect of oblique image acquisition on the accuracy of QSM and a robust tilt correction method. Magn. Reson. Med. 89, 1791–1808. 10.1002/mrm.29550

Körzdörfer, G., Kirsch, R., Liu, K., Pfeuffer, J., Hensel, B., Jiang, Y., Ma, D., Gratz, M., Bär, P., Bogner, W., Springer, E., Cardoso, P.L., Umutlu, L., Trattnig, S., Griswold, M., Gulani, V., Nittka, M., 2019. Reproducibility and Repeatability of MR Fingerprinting Relaxometry in the Human Brain. Radiology 292, 429–437. 10.1148/radiol.2019182360

Kühne, F., Neumann, W.-J., Hofmann, P., Marques, J., Kaindl, A.M., Tietze, A., 2021. Assessment of myelination in infants and young children by T1 relaxation time measurements using the magnetization-prepared 2 rapid acquisition gradient echoes sequence. Pediatr. Radiol. 51, 2058–2068. 10.1007/s00247-021-05109-5

Lai, K.-W., Aggarwal, M., van Zijl, P., Li, X., Sulam, J., 2020. Medical Image Computing and Computer Assisted Intervention – MICCAI 2020, 23rd International Conference, Lima, Peru, October 4–8, 2020, Proceedings, Part II. Lect Notes Comput Sc 12262, 125–135. 10.1007/978-3-030-59713-9_13

Langkammer, C., Pirpamer, L., Seiler, S., Deistung, A., Schweser, F., Franthal, S., Homayoon, N., Katschnig-Winter, P., Koegl-Wallner, M., Pendl, T., Stoegerer, E.M., Wenzel, K., Fazekas, F., Ropele, S., Reichenbach, J.R., Schmidt, R., Schwingenschuh, P., 2016. Quantitative Susceptibility Mapping in Parkinson’s Disease. Plos One 11, e0162460. 10.1371/journal.pone.0162460

Langkammer, C., Schweser, F., Krebs, N., Deistung, A., Goessler, W., Scheurer, E., Sommer, K., Reishofer, G., Yen, K., Fazekas, F., Ropele, S., Reichenbach, J.R., 2012. Quantitative susceptibility mapping (QSM) as a means to measure brain iron? A post mortem validation study. Neuroimage 62, 1593–1599. 10.1016/j.neuroimage.2012.05.049

Leutritz, T., Seif, M., Helms, G., Samson, R.S., Curt, A., Freund, P., Weiskopf, N., 2020. Multiparameter mapping of relaxation (R1, R2*), proton density and magnetization transfer saturation at 3 T: A multicenter dual-vendor reproducibility and repeatability study. Hum. Brain Mapp. 41, 4232–4247. 10.1002/hbm.25122

Li, G., Tong, R., Zhang, M., Gillen, K.M., Jiang, W., Du, Y., Wang, Y., Li, J., 2023. Age-dependent changes in brain iron deposition and volume in deep gray matter nuclei using quantitative susceptibility mapping. NeuroImage 269, 119923. 10.1016/j.neuroimage.2023.119923

Li, G., Zhai, G., Zhao, X., An, H., Spincemaille, P., Gillen, K.M., Ku, Y., Wang, Y., Huang, D., Li, J., 2019. 3D texture analyses within the substantia nigra of Parkinson’s disease patients on quantitative susceptibility maps and R2∗ maps. Neuroimage 188, 465–472. 10.1016/j.neuroimage.2018.12.041

Li, J., Chang, S., Liu, T., Jiang, H., Dong, F., Pei, M., Wang, Q., Wang, Y., 2015. Phase-corrected bipolar gradients in multi-echo gradient-echo sequences for quantitative susceptibility mapping. Magma (New York, N.Y.) 28, 347–355. 10.1007/s10334-014-0470-3

Li, W., Wu, B., Batrachenko, A., Bancroft-Wu, V., Morey, R.A., Shashi, V., Langkammer, C., Bellis, M.D., Ropele, S., Song, A.W., Liu, C., 2014. Differential developmental trajectories of magnetic susceptibility in human brain gray and white matter over the lifespan. Hum. Brain Mapp. 35, 2698–2713. 10.1002/hbm.22360

Li, W., Wu, B., Liu, C., 2011. Quantitative susceptibility mapping of human brain reflects spatial variation in tissue composition. Neuroimage 55, 1645–1656. 10.1016/j.neuroimage.2010.11.088

Li, Y., Sethi, S.K., Zhang, C., Miao, Y., Yerramsetty, K.K., Palutla, V.K., Gharabaghi, S., Wang, C., He, N., Cheng, J., Yan, F., Haacke, E.M., 2021. Iron Content in Deep Gray Matter as a Function of Age Using Quantitative Susceptibility Mapping: A Multicenter Study. Frontiers Neurosci. 14, 607705. 10.3389/fnins.2020.607705

Lin, P.-Y., Chao, T.-C., Wu, M.-L., 2015. Quantitative Susceptibility Mapping of Human Brain at 3T: A Multisite Reproducibility Study. Am. J. Neuroradiol. 36, 467–474. 10.3174/ajnr.a4137

Liu, C., Li, W., Johnson, G.A., Wu, B., 2011. High-field (9.4 T) MRI of brain dysmyelination by quantitative mapping of magnetic susceptibility. Neuroimage 56, 930–938. 10.1016/j.neuroimage.2011.02.024

Lodygensky, G.A., Marques, J.P., Maddage, R., Perroud, E., Sizonenko, S.V., Hüppi, P.S., Gruetter, R., 2012. In vivo assessment of myelination by phase imaging at high magnetic field. NeuroImage 59, 1979–1987. 10.1016/j.neuroimage.2011.09.057

Luo, J., He, X., Yablonskiy, D.A., 2014. Magnetic susceptibility induced white matter MR signal frequency shifts—experimental comparison between Lorentzian sphere and generalized Lorentzian approaches. Magnetic Resonance in Medicine 71, 1251–1263. 10.1002/mrm.24762

Lutti, A., Dick, F., Sereno, M.I., Weiskopf, N., 2014. Using high-resolution quantitative mapping of R1 as an index of cortical myelination. Neuroimage 93, 176–188. 10.1016/j.neuroimage.2013.06.005

Ma, D., Gulani, V., Seiberlich, N., Liu, K., Sunshine, J.L., Duerk, J.L., Griswold, M.A., 2013. Magnetic resonance fingerprinting. Nature 495, 187–192. 10.1038/nature11971

Marquand, A.F., Rezek, I., Buitelaar, J., Beckmann, C.F., 2016a. Understanding Heterogeneity in Clinical Cohorts Using Normative Models: Beyond Case-Control Studies. Biol. Psychiatry 80, 552–561. 10.1016/j.biopsych.2015.12.023

Marquand, A.F., Wolfers, T., Mennes, M., Buitelaar, J., Beckmann, C.F., 2016b. Beyond Lumping and Splitting: A Review of Computational Approaches for Stratifying Psychiatric Disorders. Biol. Psychiatry: Cogn. Neurosci. Neuroimaging 1, 433–447. 10.1016/j.bpsc.2016.04.002

Marques, J.P., Gruetter, R., 2013. New Developments and Applications of the MP2RAGE Sequence - Focusing the Contrast and High Spatial Resolution R1 Mapping. PLoS ONE 8, e69294. 10.1371/journal.pone.0069294

Marques, J.P., Kober, T., Krueger, G., van der Zwaag, W., de Moortele, P.F.V., Gruetter, R., 2010. MP2RAGE, a self bias-field corrected sequence for improved segmentation and T1-mapping at high field. Neuroimage 49, 1271–1281. 10.1016/j.neuroimage.2009.10.002

Miletić, S., Bazin, P.-L., Isherwood, S.J.S., Keuken, M.C., Alkemade, A., Forstmann, B.U., 2022. Charting human subcortical maturation across the adult lifespan with in vivo 7 T MRI. Neuroimage 249, 118872. 10.1016/j.neuroimage.2022.118872

Miller, K.L., Alfaro-Almagro, F., Bangerter, N.K., Thomas, D.L., Yacoub, E., Xu, J., Bartsch, A.J., Jbabdi, S., Sotiropoulos, S.N., Andersson, J.L.R., Griffanti, L., Douaud, G., Okell, T.W., Weale, P., Dragonu, I., Garratt, S., Hudson, S., Collins, R., Jenkinson, M., Matthews, P.M., Smith, S.M., 2016. Multimodal population brain imaging in the UK Biobank prospective epidemiological study. Nat. Neurosci. 19, 1523–1536. 10.1038/nn.4393

Najdenovska, E., Alemán-Gómez, Y., Battistella, G., Descoteaux, M., Hagmann, P., Jacquemont, S., Maeder, P., Thiran, J.-P., Fornari, E., Cuadra, M.B., 2018. In-vivo probabilistic atlas of human thalamic nuclei based on diffusion-weighted magnetic resonance imaging. Sci. Data 5, 180270. 10.1038/sdata.2018.270

Oldehinkel, M., Llera, A., Faber, M., Huertas, I., Buitelaar, J.K., Bloem, B.R., Marquand, A.F., Helmich, R.C., Haak, K.V., Beckmann, C.F., 2022. Mapping dopaminergic projections in the human brain with resting-state fMRI. Elife 11, e71846. 10.7554/elife.71846

Park, H.-J., Westin, C.-F., Kubicki, M., Maier, S.E., Niznikiewicz, M., Baer, A., Frumin, M., Kikinis, R., Jolesz, F.A., McCarley, R.W., Shenton, M.E., 2004. White matter hemisphere asymmetries in healthy subjects and in schizophrenia: a diffusion tensor MRI study. NeuroImage 23, 213–223. 10.1016/j.neuroimage.2004.04.036

Penny, W.D., Friston, K.J., Ashburner, J.T., Kiebel, S.J., Nichols, T.E. (Eds.), 2006. Statistical Parametric Mapping: The Analysis of Functional Brain Images. Elsevier.

Persson, N., Wu, J., Zhang, Q., Liu, Ting, Shen, J., Bao, R., Ni, M., Liu, Tian, Wang, Y., Spincemaille, P., 2015. Age and sex related differences in subcortical brain iron concentrations among healthy adults. Neuroimage 122, 385–398. 10.1016/j.neuroimage.2015.07.050

Piredda, G.F., Caneschi, S., Hilbert, T., Bonanno, G., Joseph, A., Egger, K., Peter, J., Klöppel, S., Jehli, E., Grieder, M., Slotboom, J., Seiffge, D., Goeldlin, M., Hoepner, R., Willems, T., Vulliemoz, S., Seeck, M., Venkategowda, P.B., Jerez, R.A.C., Maréchal, B., Thiran, J., Wiest, R., Kober, T., Radojewski, P., 2023. Submillimeter T1 atlas for subject-specific abnormality detection at 7T. Magn. Reson. Med. 89, 1601–1616. 10.1002/mrm.29540

Piredda, G.F., Hilbert, T., Granziera, C., Bonnier, G., Meuli, R., Molinari, F., Thiran, J., Kober, T., 2020. Quantitative brain relaxation atlases for personalized detection and characterization of brain pathology. Magn. Reson. Med. 83, 337–351. 10.1002/mrm.27927

Pomponio, R., Erus, G., Habes, M., Doshi, J., Srinivasan, D., Mamourian, E., Bashyam, V., Nasrallah, I.M., Satterthwaite, T.D., Fan, Y., Launer, L.J., Masters, C.L., Maruff, P., Zhuo, C., Völzke, H., Johnson, S.C., Fripp, J., Koutsouleris, N., Wolf, D.H., Gur, Raquel, Gur, Ruben, Morris, J., Albert, M.S., Grabe, H.J., Resnick, S.M., Bryan, R.N., Wolk, D.A., Shinohara, R.T., Shou, H., Davatzikos, C., 2020. Harmonization of large MRI datasets for the analysis of brain imaging patterns throughout the lifespan. NeuroImage 208, 116450. 10.1016/j.neuroimage.2019.116450

Pontillo, G., Petracca, M., Monti, S., Quarantelli, M., Lanzillo, R., Costabile, T., Carotenuto, A., Tortora, F., Elefante, A., Morra, V.B., Brunetti, A., Palma, G., Cocozza, S., 2022. Clinical correlates of R1 relaxometry and magnetic susceptibility changes in multiple sclerosis: a multi-parameter quantitative MRI study of brain iron and myelin. Eur Radiol 1–10. 10.1007/s00330-022-09154-y

Pyatigorskaya, N., Sanz-Morère, C.B., Gaurav, R., Biondetti, E., Valabregue, R., Santin, M., Yahia-Cherif, L., Lehéricy, S., 2020. Iron Imaging as a Diagnostic Tool for Parkinson’s Disease: A Systematic Review and Meta-Analysis. Front. Neurol. 11, 366. 10.3389/fneur.2020.00366

QSM Consensus Organization Committee, Bilgic, B., Costagli, M., Chan, K., Duyn, J., Langkammer, C., Lee, J., Li, X., Liu, C., Marques, J.P., Milovic, C., Robinson, S.D., Schweser, F., Shmueli, K., Spincemaille, P., Straub, S., van Zijl, P., Wang, Y., ISMRM of quantitative susceptibility mapping for clinical research in the brain: A consensus of the ISMRM electro-magnetic tissue properties study group. Magn. Reson. Med. 91, 1834– 1862. 10.1002/mrm.30006

Ramanoël, S., Hoyau, E., Kauffmann, L., Renard, F., Pichat, C., Boudiaf, N., Krainik, A., Jaillard, A., Baciu, M., 2018. Gray Matter Volume and Cognitive Performance During Normal Aging. A Voxel-Based Morphometry Study. Front Aging Neurosci 10, 235. 10.3389/fnagi.2018.00235

Rooney, W.D., Johnson, G., Li, X., Cohen, E.R., Kim, S.-G., Ugurbil, K., Springer, C.S., 2007. Magnetic field and tissue dependencies of human brain longitudinal 1H2O relaxation in vivo. Magnetic resonance in medicine 57, 308–318. 10.1002/mrm.21122

Rutherford, S., Barkema, P., Tso, I.F., Sripada, C., Beckmann, C.F., Ruhe, H.G., Marquand, A.F., 2023. Evidence for embracing normative modeling. eLife 12, e85082. 10.7554/elife.85082

Rutherford, S., Fraza, C., Dinga, R., Kia, S.M., Wolfers, T., Zabihi, M., Berthet, P., Worker, A., Verdi, S., Andrews, D., Han, L.K., Bayer, J.M., Dazzan, P., McGuire, P., Mocking, R.T., Schene, A., Sripada, C., Tso, I.F., Duval, E.R., Chang, S.-E., Penninx, B.W., Heitzeg, M.M., Burt, S.A., Hyde, L.W., Amaral, D., Nordahl, C.W., Andreasssen, O.A., Westlye, L.T., Zahn, R., Ruhe, H.G., Beckmann, C., Marquand, A.F., 2022a. Charting brain growth and aging at high spatial precision. eLife 11, e72904. 10.7554/elife.72904

Rutherford, S., Kia, S.M., Wolfers, T., Fraza, C., Zabihi, M., Dinga, R., Berthet, P., Worker, A., Verdi, S., Ruhe, H.G., Beckmann, C.F., Marquand, A.F., 2022b. The normative modeling framework for computational psychiatry. Nat Protoc 17, 1711–1734. 10.1038/s41596-022-00696-5

Satterthwaite, T.D., Connolly, J.J., Ruparel, K., Calkins, M.E., Jackson, C., Elliott, M.A., Roalf, D.R., Hopson, R., Prabhakaran, K., Behr, M., Qiu, H., Mentch, F.D., Chiavacci, R., Sleiman, P.M.A., Gur, R.C., Hakonarson, H., Gur, R.E., 2016. The Philadelphia Neurodevelopmental Cohort: A publicly available resource for the study of normal and abnormal brain development in youth. NeuroImage 124, 1115–1119. 10.1016/j.neuroimage.2015.03.056

Scahill, R.I., Frost, C., Jenkins, R., Whitwell, J.L., Rossor, M.N., Fox, N.C., 2003. A Longitudinal Study of Brain Volume Changes in Normal Aging Using Serial Registered Magnetic Resonance Imaging. Arch Neurol-chicago 60, 989–994. 10.1001/archneur.60.7.989

Seiler, A., Schöngrundner, S., Stock, B., Nöth, U., Hattingen, E., Steinmetz, H., Klein, J.C., Baudrexel, S., Wagner, M., Deichmann, R., Gracien, R.-M., 2020. Cortical aging – new insights with multiparametric quantitative MRI. Aging Albany Ny 12, 16195–16210. 10.18632/aging.103629

Snoek, L., van der Miesen, M.M., Beemsterboer, T., van der Leij, A., Eigenhuis, A., Scholte, H.S., 2021. The Amsterdam Open MRI Collection, a set of multimodal MRI datasets for individual difference analyses. Sci. Data 8, 85. 10.1038/s41597-021-00870-6

Stüber, C., Morawski, M., Schäfer, A., Labadie, C., Wähnert, M., Leuze, C., Streicher, M., Barapatre, N., Reimann, K., Geyer, S., Spemann, D., Turner, R., 2014. Myelin and iron concentration in the human brain: A quantitative study of MRI contrast. Neuroimage 93, 95–106. 10.1016/j.neuroimage.2014.02.026

Taki, Y., Thyreau, B., Kinomura, S., Sato, K., Goto, R., Kawashima, R., Fukuda, H., 2011. Correlations among Brain Gray Matter Volumes, Age, Gender, and Hemisphere in Healthy Individuals. PLoS ONE 6, e22734. 10.1371/journal.pone.0022734

Teixeira, R.P.A.G., Neji, R., Wood, T.C., Baburamani, A.A., Malik, S.J., Hajnal, J.V., 2019. Controlled saturation magnetization transfer for reproducible multivendor variable flip angle T1 and T2 mapping. Magnetic resonance in medicine mrm.28109. 10.1002/mrm.28109

Terribilli, D., Schaufelberger, M.S., Duran, F.L.S., Zanetti, M.V., Curiati, P.K., Menezes, P.R., Scazufca, M., Amaro, E., Leite, C.C., Busatto, G.F., 2011. Age-related gray matter volume changes in the brain during non-elderly adulthood. Neurobiol Aging 32, 354–368. 10.1016/j.neurobiolaging.2009.02.008

Treit, S., Naji, N., Seres, P., Rickard, J., Stolz, E., Wilman, A.H., Beaulieu, C., 2021. R2* and quantitative susceptibility mapping in deep gray matter of 498 healthy controls from 5 to 90 years. Hum. Brain Mapp. 42, 4597–4610. 10.1002/hbm.25569

Veraart, J., Novikov, D.S., Christiaens, D., Ades-aron, B., Sijbers, J., Fieremans, E., 2016. Denoising of diffusion MRI using random matrix theory. Neuroimage 142, 394–406. 10.1016/j.neuroimage.2016.08.016

Watkins, K.E., Paus, T., Lerch, J.P., Zijdenbos, A., Collins, D.L., Neelin, P., Taylor, J., Worsley, K.J., Evans, A.C., 2001. Structural Asymmetries in the Human Brain: a Voxel-based Statistical Analysis of 142 MRI Scans. Cereb. Cortex 11, 868–877. 10.1093/cercor/11.9.868

Wharton, S., Bowtell, R., 2015. Effects of white matter microstructure on phase and susceptibility maps. Magnetic Resonance in Medicine 73, 1258–1269. 10.1002/mrm.25189

Xu, J., Kobayashi, S., Yamaguchi, S., Iijima, K., Okada, K., Yamashita, K., 2000. Gender effects on age-related changes in brain structure. AJNR Am. J. Neuroradiol. 21, 112–8.

Yeatman, J.D., Wandell, B.A., Mezer, A.A., 2014. Lifespan maturation and degeneration of human brain white matter. Nature communications 5, 4932. 10.1038/ncomms5932

Zhang, Y., Wei, H., Cronin, M.J., He, N., Yan, F., Liu, C., 2018. Longitudinal atlas for normative human brain development and aging over the lifespan using quantitative susceptibility mapping. Neuroimage 171, 176–189. 10.1016/j.neuroimage.2018.01.008

