## Supplementary for "Normative trajectories of R_1_, R_2_* and magnetic susceptibility in basal ganglia on healthy ageing"

### Supplementary Materials

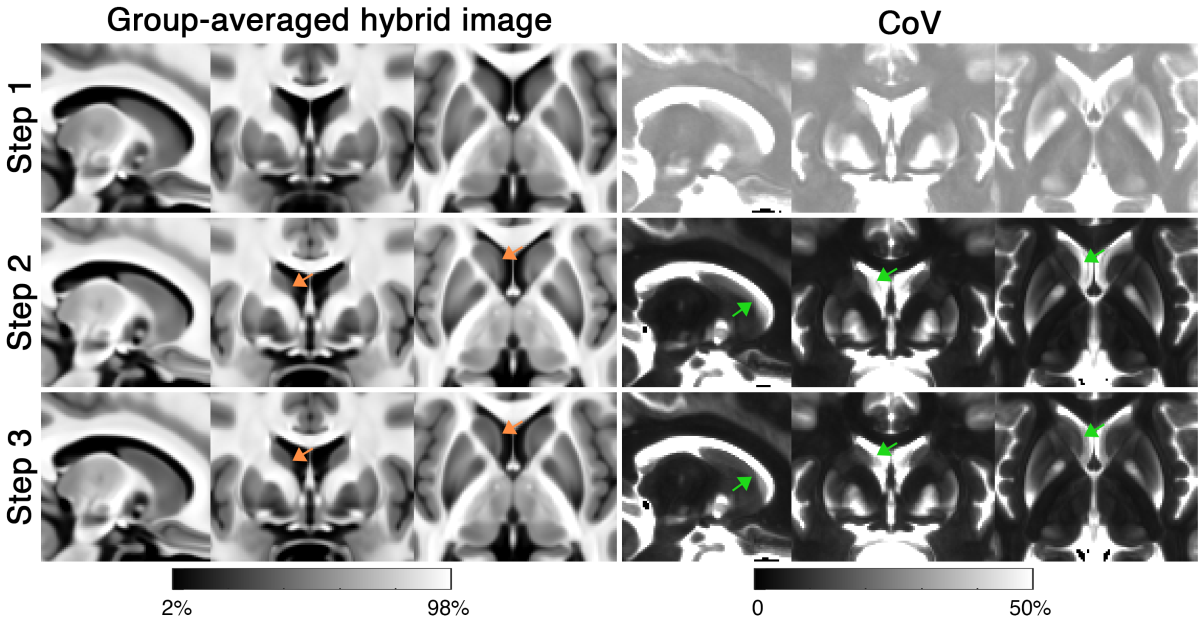


Figure S1: (left) Group-averaged hybrid image derived using the results of each registration step and (right) the corresponding coefficient of variation (CoV) image across all subjects. The result of Step 1 shows that the contrast between the globus pallidus and putamen is low without matching the contrast as in Step 2, resulting in blurry edges between the two close proximal structures. Note the boundary between the caudate nucleus and ventricle is notably sharper in the hybrid image of Step 3 compared to Step 2 (orange arrows). The CoV images also show lower values in the caudate nucleus/ventricle boundary at Step 3 (green arrows).

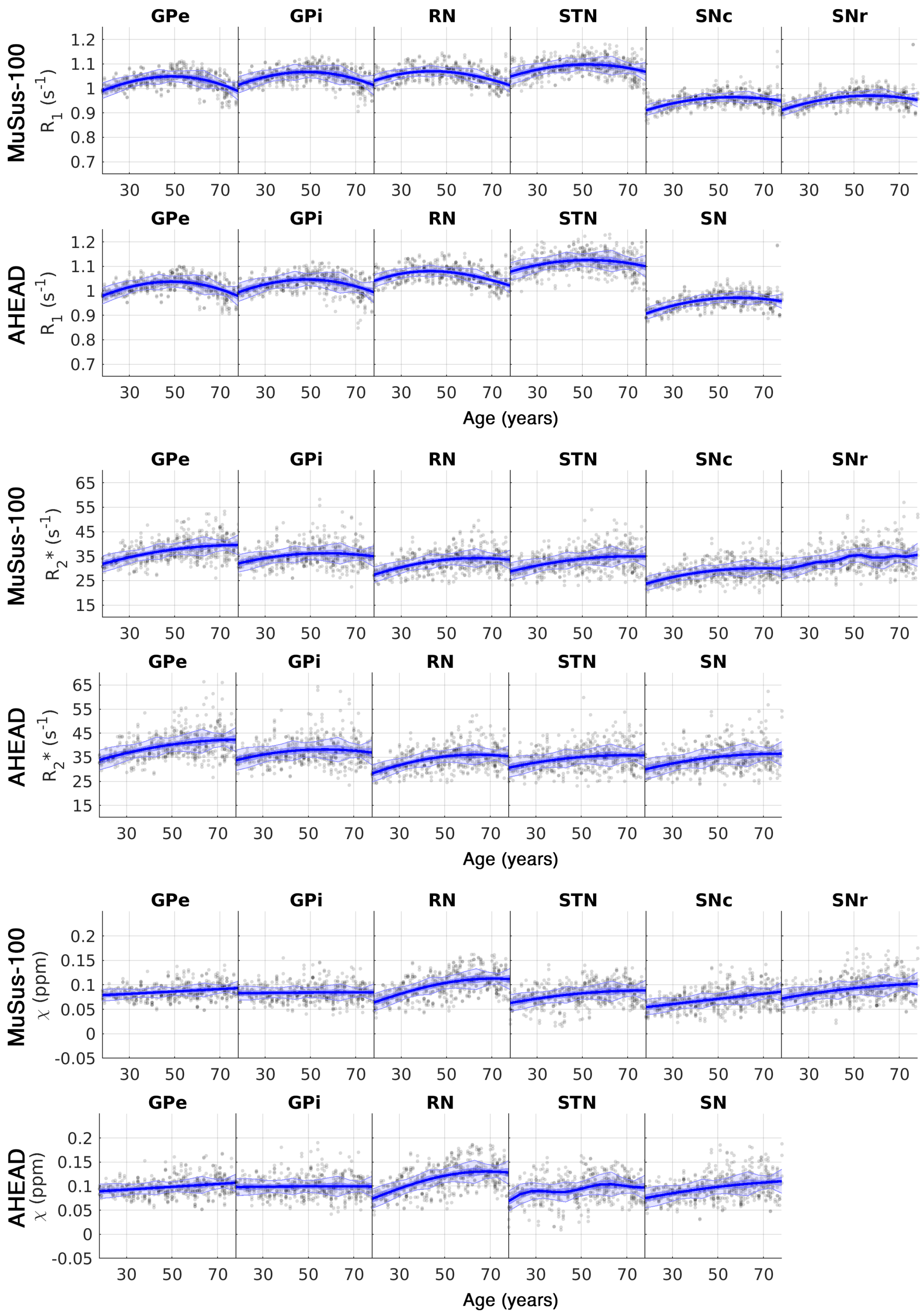


Figure S2: Normative trajectories derived using the AHEAD atlas (Alkemade et al., 2020) following the same process as described in the Methods section. The basal ganglia labels were obtained by hard thresholding the probabilistic images at 75%. The trajectories of the same structure derived using the MuSus-100 atlas are displayed alongside with the AHEAD atlas results for direct comparison.

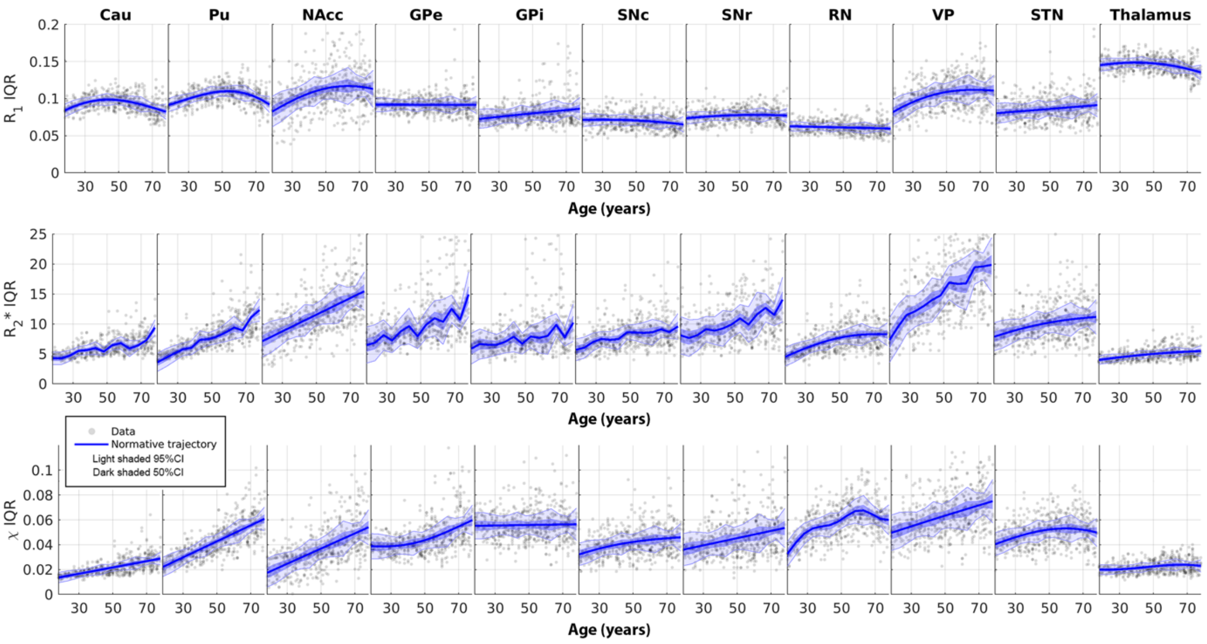


Figure S3: Normative trajectories of the interquartile range (IQR) of $R_1$, $R_2$*, $\chi$ as a function of age for 11 different deep grey matter structures present in the MuSus-100 atlas. The data points shown on the scatter plot are corrected for sex and hemispheric effects.

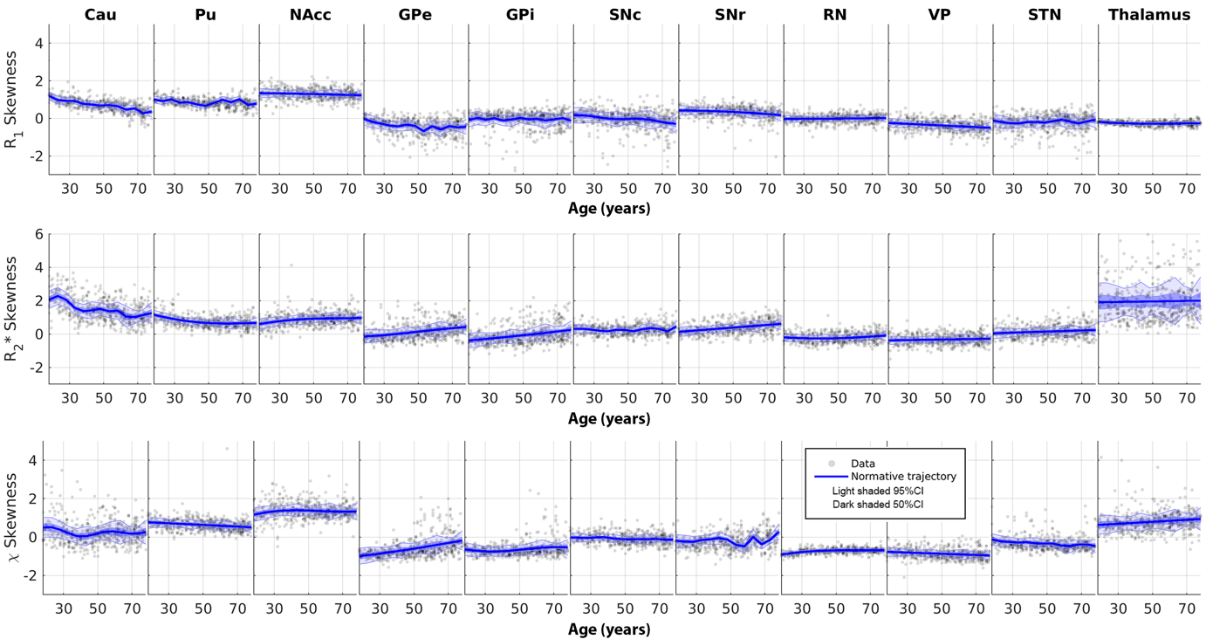


Figure S4: Normative trajectories of the skewness of the distribution of $R_1$, $R_2$*, $\chi$ as a function of age for 11 different deep grey matter structures present in the MuSus-100 atlas. The data points shown on the scatter plot are corrected for sex and hemispheric effects.

### Reference


Alkemade, A., Mulder, M.J., Groot, J.M., Isaacs, B.R., Berendonk, N. van, Lute, N., Isherwood, S.J., Bazin, P.-L., Forstmann, B.U., 2020. The Amsterdam Ultra-high field adult lifespan database (AHEAD): A freely available multimodal 7 Tesla submillimeter magnetic resonance imaging database. Neuroimage 221, 117200. https://doi.org/10.1016/j.neuroimage.2020.117200
